# Relative Reduction of the Biological and phylogenetic diversity of oral microbiome in diabetic and pre-diabetic subjects

**DOI:** 10.1101/406736

**Authors:** Amr T. M. Saeb, Khalid A. Al-Rubeaan, Khalid Aldosary, G. K. Udaya Raja, Balavenkatesh Mani, Mohamed Abouelhoda, Hamsa T. Tayeb

## Abstract

**Background:** There is a suggested reciprocal relationship between oral health and systemic disease such as type 2 diabetes. In this relationship, a systemic disease predisposing to oral infection, and when that infection is present, the oral infection aggravates the progression of the systemic disease. Several studies suggested that some oral microbiome constituents are linked to both diabetes, metabolic syndrome and obesity. This study aims to compare the microbial diversity and population structure of oral microbiome among normoglycemic, impaired glucose tolerance (IGT) and diabetic subjects.

**Methodology:** This study followed a case-control design (15 T2D patients, 10 IGTs and, 19 controls). Patient records were screened as per the inclusion and exclusion criteria. Assessment of periodontitis and oral health was performed to all subjects. DNA Isolation purification and quantification from collected Saliva samples were performed. 16SrRNA hypervariable regions were amplified and sequenced. Generated sequences were subjected to bioinformatics analysis. Statistical analysis and diversity indices were computed with the statistical software R, the vegan R-package, and Past318 software.

**Results:** A total observed number of 551 OTUs. A clear reduction of the number of species (OTUs) was observed in both IGT (412) and diabetic group (372) compared with the normoglycemic group (502). This was associated with a similar pattern of biological diversity among the three groups. Phylogenetic diversity (PD-SBL) value in the normoglycemic group was higher than the diabetic group. The diabetic group had the highest evenness value and the highest microbiome bacterial pathogenic content.

**Conclusion:** We observed a clear reduction in the biological and phylogenetic diversity in the diabetic and pre-diabetic oral microbiome in comparison with the normoglycemic oral microbiome. However, this reduction was associated with an increase in the pathogenic content of the hyperglycemic microbiomes.

## Background

Diabetes is the leading cause of morbidity and mortality all over the world. Type 2 diabetes prevalence was 415 million people in 2015 and projected to reach 642 million by 2040 [1]. Diabetes is known as a significant risk factor for rigorous and progressive periodontitis, infection or lesions resulting in the obliteration of tissues and supporting bone that form the attachment around a tooth. Whereas the oral cavity presents a permanent source of infectious agents, and its state often mirrors the progression of systemic pathologies. Diabetes and periodontitis are thought to share a common pathogenesis that involves triggering inflammatory responses that can be perceived at both local and systemic levels. This reciprocal relationship between diabetes and oral health has been supported by lots of evidence [2]. The reciprocal relationship presents an example of systemic disease predisposing to oral infection, and when that infection is present, the oral infection aggravates the progression of the systemic disease. In addition, several studies suggested that controlling one arm in this reciprocal relationship can lead to improvement of the status of the other arm [2–4]. It was proposed that the increase of advanced glycation end products (AGEs) because of the chronic hyperglycemic state accompanied by the presence of infection and an embellished host response may provide a possible justification for the clinical outcomes observed in diabetic patients with periodontal disease. Furthermore, bacterial cellular components and products such as lipopolysaccharide (LPS) and endotoxin as well play a role in aggravating an inflammatory response in the host cells through the Toll-like protein receptors (TLRs) and thus can induce an inflammatory cascade [5,6]. Thus, the oral infection can be an indicator for incidence and progression of diabetes. The inflammatory response is essentially caused by the chronic effects of hyperglycemia and specifically the presence of biological reactive glycated proteins and lipids that induce inflammatory responses. In turn, the enhanced innate immune responses and periodontal tissue destruction related to an altered inflammatory response. Since periodontitis can be more than just a localized oral infection, the effects have been hypothesized to be systematic [2,5]. Although little studied, the oral microbiome may be important in chronic diseases as diabetes through direct metabolism of chemical carcinogens and through systemic inflammatory effects [7–9]. Recently, it was suggested that diabetes-enhanced IL-17 alters the oral microbiota and renders it more pathogenic [10]. Oral microbiome can affect systemic health through inhibition of potential pathogen, regulation of immune responses, nutrients absorption and metabolism [11]. In addition, oral microbiome bacteria and/or its by-products can traffic through the human body and lead to systemic diseases such as cardiovascular disease, Alzheimer’s disease and cancers [9,12–15]. Several studies suggested that some oral microbiome constituents are linked to both Diabetes, metabolic syndrome and obesity. For example, the presence of *Porphyromonas gingivalis,* especially clones with type II fimbriae, in periodontal pockets effects the glycemic level in diabetes [16]. Moreover, poor glycemic control was associated with increased cell numbers of red complex bacteria (*Porphyromonas gingivalis*, *Tannerella forsythia*, and *Treponema denticola*) in subgingival biofilm [17]. In addition, though *Actinobacillus actinomycetemcomitans*, *Campylobacter rectus*, *Capnocytophaga spp*, *Eikenella corrodens*, *Fusobacterium nucleatum*, *Porphyromonas gingivalis*, *Prevotella intermedia* were observed in both diabetics and non-diabetics., *Porphyromonas gingivalis* was found significantly more among diabetics [18]. Furthermore, it was found that diabetic had slightly higher levels of oral bacteria compared with non-diabetics. Moreover, subjects with HbA1c > 6.5% had rather lower levels of Bifidobacteria in the mouth and stool [19]. In addition, it was found that the numbers of total streptococci and lactobacilli were significantly higher in supragingival plaque from diabetics than in nondiabetics. Furthermore, Lactobacillus bacterial numbers were elevated in diabetics with active caries, although salivary eubacterial DNA profiles were not significantly altered [20]. Most recently, Long et al., showed that the phylum Actinobacteria was present significantly less abundant among patients with diabetes than among the controls. Moreover, multiple bacteria taxa in the phylum Actinobacteria are associated with the risk of type 2 diabetes. However, their significant study limitation was that they only classify the oral bacteria at only the genus level in relation to diabetes risk [21]. In this study aims to compare the microbial diversity and population structure of oral microbes among normoglycemic, glucose intolerant and diabetic patients using a highly deep sequencing technique to the species level. In addition, it also aimed to the identification of the bacterial species combination that are can be used as early biomarkers of diabetes.

## Materials and Methods

This is a cross-sectional case-control hospital-based study involving three cohorts; namely normoglycemic, pre-diabetic and type 2 diabetic subjects. They were recruited during the period from May 2013 to March 2015 from the University Diabetes Center, King Saud University. The total recruited subjects were 44 subjects divided into 19 normoglycemic subjects used as a control, 10 impaired glucose tolerant subjects known as pre-diabetic and 15 type 2 diabetic patients. Subjects included in this study were any gender aged between 40 and 55 years and Saudi nationals. Any subjects used an antibiotic for the last 6 months or known to have severe periodontal disease was excluded. Immunocompromised patients like subjected with post organ transplant, end-stage renal disease (ESRD), severely debilitating diseases: cancer or receiving chemotherapy were not included in this study. This study did not include patients with autoimmune diseases or pregnant women. For the diabetic cohort, any patients with type 1, gestational and other types of diabetes were excluded. Normal subjects and pre-diabetic subjects were recruited from patients’ relatives attending the clinics after being subjected to 100 gm oral glucose tolerance (OGT) test after 10 hours of the overnight fast. The American Diabetes Association (ADA) criteria were used, where normal subjects had FPG less than 100 mg/dl and or, 2-hour post-load glucose less than 140 mg/dl and prediabetic subjects who had FPG 100-125 mg/dl, 2-hour post-load glucose between 140 and 199 mg/dl. The type 2 diabetic patients were recruited from patients’ cohort attending University Diabetes center clinics diagnosed ADA criteria of (FPG) ≥126 mg/dl or Two-hour plasma glucose ≥200 mg/dl and managed with oral hypoglycemic agents only.

Each subject was interviewed individually by a research physician and was consented. General demographic data including age, gender, weight, height and history of any dental problems, in addition to drug history and mouth hygiene practice. For diabetic patients, extra data were collected including diabetes duration, the presence of chronic complications or associated disease. The degree of metabolic control was collected from the patient file including HbA1c, FPG and 2 hours post-prandial.

### Periodontal assessment

All the 44 subjects were referred to a specialized dental clinic for periodontal assessment. A full mouth clinical examination was conducted by three examiners who had been previously exposed to the National Health and Nutrition Examination Survey (NHANES) reference examiner (Dr. Bruce Dye). The Periodontal disease was assessed by measuring the probing pocket depth (PPD) and clinical attachment loss (CAL) at six sites involving: distobuccal, mid-buccal, mesiobuccal, distolingual, mid-lingual, and mesiolingual buccal for all teeth excluding the third molars. Measurements were taken using a periodontal probe (product number PCP2; Hu-Friedy, Chicago, IL, USA) and rounded off upwards to the nearest millimeter. Periodontitis was defined according to the Centers for Disease Control and Prevention/American Academy of Periodontology (CDC/AAP) [22]. Severe periodontitis was defined as having at least two interproximal sites with CAL ≥ 6 mm (not on the same tooth) and at least one interproximal site with PPD ≥ 5 mm. Moderate periodontitis was defined as having at least two interproximal sites with CAL ≥ 4 mm (not on the same tooth) or at least two interproximal sites with PPD ≥ 5 mm (not on the same tooth). Mild periodontitis was defined as having at least two interproximal sites with CAL ≥ 3 mm and at least two interproximal sites with PPD ≥ 4 mm (not on the same tooth) or one site with PPD ≥ 5 mm. During the measurement of PPD, a periodontal probe was inserted to the base of the sulcus or pocket with a maximum force of 20 g and bleeding on probing (BOP) was positive if the probed site bled about 20 seconds after probing the lingual and buccal surfaces of each tooth. The BOP was classified as high if 30% or more teeth showed positive BOP, and as low otherwise [23]. The Silness and Loe Plaque Index, a measure of oral hygiene status, was determined by visual assessment of the presence of bacterial plaque after passing a periodontal probe around the tooth surface of six pre-selected Ramfjford teeth. The plaque index was coded as 0 if no plaque was present, 1 if the dental plaque was present after passing the periodontal probe around the tooth, 2 if the plaque was visible along the gingival margin, and 3 if the tooth surface was covered with abundant plaque.

### Molecular Analysis

All participants in this study were asked to provide saliva in the Omingene saliva collection tubes of Kit OM501 for microbial DNA analysis following the instructions of the manufacturer after rubbing tong around for about 60 seconds. Prior to sample collection, subjects were instructed to abstain from eating, drinking, smoking, and mouth washing for at least 60 min. The sample was placed immediately in an icebox and was transferred to the strategic center for diabetes research laboratory, were they stored at – 20C till DNA isolation and purification.

### Bacterial DNA Isolation purification, quantification and enrichment

All saliva samples collected from the studied subjects were used to isolate bacterial DNA using Promega Maxwell 16 automated DNA isolation machine according to the instructions of the manufacturer. The isolated DNA samples were quantified using the NanoDrop 2000c UV-Vis spectrophotometer. Agilent 2100 Bioanalyzer system was used for sizing, quantitation and quality control assessment for the isolated DNA. The NEBNext^®^ Microbiome DNA was used to eliminate human DNA and to facilitate enrichment of microbial DNA in the samples. This kit selectively binds and remove the CpG-methylated host DNA to keep microbial diversity intact after enrichment. All the isolated DNA samples were then stored at −20 C.

### 16SrRNA gene hypervariable regions amplification

16S rRNA gene was amplified using Ion 16S™ Metagenomics Kit (Thermo Fisher Scientific) following the instructions of the manufacturer. To increase the resolving power of 16S rRNA profiling, the primers were designed to amplify variable regions 2, 4, and 8 in a single tube with the resulting amplicon fragments of ∼250 base pairs (bp), ∼288 bp and ∼295 bp, respectively. In a second single tube, a multiplex PCR reaction targets variable regions 3, 6-7 and 9 with resulting amplicon fragments of ∼215 bp, ∼260 bp and ∼209 bp, respectively. The primer pools in the Kit are designed to target >80% of sequences found in the Greengenes database with 100% identity for a primer pair amplifying at least one variable region. The PCR amplifications were performed using Applied Biosystems VeritiTM 96-Well Thermal Cycler under the following cycling conditions: 95°C for 10 min, then Cycle 18–25 cycles 95°C for 30 sec, 58°C for 30 sec and 72°C for 20 sec. Followed by a final extension step at 72°C for 7 min then Hold at 4°C overnight. The amplicons were purified using an Agencourt AMPure XP Kit (Beckman Coulter, Brea, CA, USA) according to the manufacturer’s instructions. The PCR amplified products were used for library preparation using Ion plus Fragment Library kit with sample indexing using IonXpress Barcode adapters 1-16 kit following the manufacturer’s instructions. The Library concentration was measured using the Ion universal library quantification kit and diluted to 13 pM. The Diluted libraries were equally pooled together and used as a template for emulsion PCR. Emulsion PCR and enrichment of template-positive particles were performed using an Ion PGM Hi-Q OT2 Kit (Thermo Fisher Scientific) and the Ion OneTouch 2 system (Life Technologies) according to the manufacturer’s instructions. The enriched template-positive Ion sphere particles were loaded onto an Ion 318 chip v2 (Thermo Fisher Scientific), and sequenced on an Ion PGM (Thermo Fisher Scientific) using an Ion PGM Hi-Q Sequencing Kit (Thermo Fisher Scientific) according to the manufacturer’s instructions to ensure high degree of sensitivity required and the deal with high complexity of our samples under investigation.

### DNA sequence analysis

Data analysis carried out with automated streamlined software’s Torrent Suite Software v5.0 (TSS) and IonReporter Software v5.0. Torrent Suite Software performed sequencing base call from PGM instrument and IonReporter used for annotation and taxonomical assignments. Standard analysis parameter used in Torrent Suite Software at primary base calling analysis including trimming low-quality 3′ ends of reads, removing duplicated reads, filtering out entire reads with average quality score less than Q20 (base call quality), removing of adapter sequence, removal of lower-quality 3′ ends with Low-Quality Scores, removing short reads, removing adapter dimers, removing polyclonal reads etc. Q20 is Phred scale score based on error probability (−10×log10). Q20 corresponds to a predicted error rate of one percent. AQ20 is read length at which the error rate is 1% or less. IonTorrent PGM generated sequences saved in binary alignment map (uBam) file format consist base call sequences, flow (flowgram), and qual (quality score). IonReporter Uploader plugin used to transfer all samples high-quality data from Torrent Suite Software to cloud Ion Reporter Software.

All three groups Normal Glycemic, Impaired Glucose Tolerance and Diabetic Group samples analyses launched in IonReporter Software with 16S Metagenomics Analysis Workflow. Metagenomics workflow contains curated Greengenes and premium curated MicroSEQ ID 16S rRNA reference databases. Greengenes databases manually custom curated from available existing public library because it is relatively large references containing more than 1.2 million references. As Greengenes is mostly aimed towards taxonomy information for family level and above, taxonomy information for genus and species levels was missing for almost 1 million references. A custom utility was used to curate Greengenes for genus and species-level information by querying NCBI. The final library contained 215,967 references directly from Greengenes, and 146,822 references with updated genus and species names from NCBI, totaling 362,789 references. Both the databases library were converted into a BLAST compatible format for the analysis. Analysis pipeline includes annotation, taxonomic assignment, and classification of reads is through alignment to either the curated MicroSEQ ID or curated Greengenes databases.

IonReporter identifies chimeric sequences based on a built-in algorithm and removes before it passing to subsequent downstream analysis. Each group samples analyzed together, reads were classified at species level with the taxonomical order with 16S Metagenomics Analysis Workflow. Analysis steps include trimming primer and length check after trimmed reads, processed reads placed in a hash table. This hash table contains all the unique reads and their abundance or copy number. When adding a new read to the hash table, the read will have a copy number of 1 if it does not exist in the table already, and if it does, the copy number will be increased by 1. After adding all reads, the hash table contained all unique reads, each with a copy number. Followed by all unique reads are converted to FASTA file format. Each read has a unique ID to keep analysis track in results throughout the process. Multistage BLAST search of reads against databases, this algorithm generates the input files and starts one BLAST process for each sample. BLAST parameters included E value equal to 0.01; result E value is larger than 0.01 was omitted in the results output. This final ion-reporter generated results consist of species-level OUTs (operational taxonomic unit) identification, also provides primer information, classification information, percent ID, and mapping information etc.

In IonReporter some of the OUTs not solved at the species level, those unsolved sequences retrieved and performed BLAST in Human Oral Microbial Databases (HOMD). For each unsolved sequence, the top four hits result used for further species level assignment based on following criteria. In the top four hits result, identities matching with 100% directly assigned species to the respective sequence. If none of the hits 100% identities, then selected maximum percentage identities with solved species. Even in HOMD, some sequences are not solved at the species level and these unsolved cases represented with a respective clone or hot strain species obtained in BLAST output. All sequence data were submitted to NCBI and submission number is SUB4304342.

### Statistical and phylogenetic analysis

All statistical and diversity indices were computed with the statistical software R, the vegan R-package and Past318 software [24]. MEGA 7 software was used to calculate phylogenetic diversity values [25].

## Results

### Subjects’ Recruitment

A total of 800 subjects were screened from patients followed at University Diabetic center or their normal companions. Out of which 180 patients had fulfilled the inclusion criteria were type 2 diabetes aged 40 to 55 years and Saudi nationals. Those patients were interviewed by research physicians and only 44 subjects fulfilled the study protocol after excluding any subjects used an antibiotic for the last 6 months or known to have a severe periodontal disease. Patients on folic acid or refusing dental appointment or providing saliva sample were excluded. The final study cohort included 19 normoglycemic subjects, 10 individuals with impaired glucose tolerance and 15 type 2 patients with diabetes. The selected study cohort had a different age range been 41-56 years, 33-53 years and 45-53 years for normal, IGT and diabetic groups respectively. Males were more predominant than females in the normal and diabetic group but were equal in IGT patients. Mild periodontitis was found in one diabetic patient, three IGT patients and four normal subjects as shown in **Table S1**. The mean diabetes duration was 11.67 years ranging from 3 to 20 years. There was one patient with diabetic nephropathy and proliferative retinopathy. HbA1c was ranging between 6.4% and 27.6% with a mean of 9.61% while fasting blood sugar values ranged between 5-18.3 mmol/L and a mean of 10.21 mmol/L. None of the medications prescribed to all diabetic subjects were having known effect on the oral microbiome and all normal subjects were not taking any medication during the study period

### Sequencing information and Taxonomic assignment of reads

We determined the bacterial compositions in the subjects in this study using 16S rRNA gene amplicon analysis with an Ion PGM. After applying the read quality and length filters, a total of 24568466 raw reads were obtained, while, the number of reads per samples ranged between 293442-4016984 with a mean of 558374 reads per sample. Though ion reporter software automatically removes chimeric reads, UCHIME, Chimera Slayer and Decipher identified only 0.66% chimeric reads that were later excluded. The average read length was 175 bases. The sequences were assigned to a total number of 551 operational taxonomic units (OTUs), using a cutoff distance of 0.05, by sequence comparison to updated-HOMD and trusted-HOMDext **(Figure 1)**.

**Figure 1:**
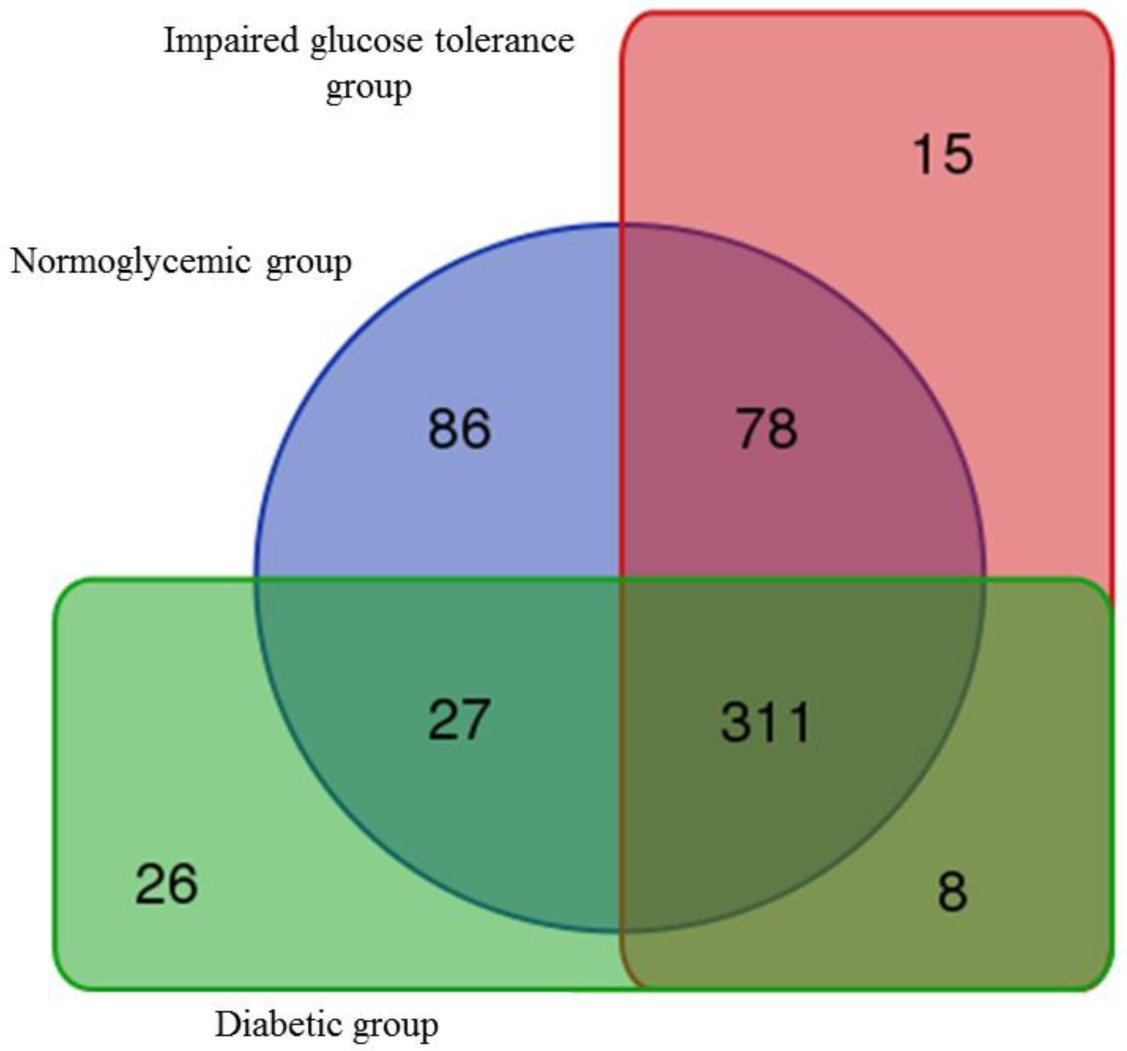
Distribution of the observed oral microbiome species (OTUs) among the different studied Glycemic groups.

Most of these reads (99%) were to the species level. The total number of species (OTUs) observed in the normoglycemic group was 502, while the number of the observed OTUs in the impaired glucose tolerance group was 412, while only 372 OTUs were observed in the diabetic group. The rarefaction curve for the number of observed species (OTUs) per sample almost reached a plateau after 10000 to 15000 sequence reads (Fig. S1). The number of shared species among the three studied groups was 311. While the number of shared (OTUs) only between the normoglycemic group and impaired glucose tolerance group was 78. Whereas the number of shared (OTUs) only between the normoglycemic group and the diabetic group was 27. The number of shared (OTUs) only between impaired glucose tolerance and the diabetic group is 8. The number of species (OTUs) observed only in the normoglycemic group was 86, while 15 species (OTUs) were observed only in the impaired glucose tolerance group and 26 species (OTUs) were observed only in the diabetic group (Table. S2).

### The core bacteriome class level

For the overall sample population (44 individuals), the number of observed classes that represents the core taxa (95% of reads) was 11 classes all belonging to domain bacteria. Namely, Actinobacteria, Bacilli, Bacteroidia, Betaproteobacteria, Clostridia, Epsilonproteobacteria, Erysipelotrichia, Flavobacteriia, Gammaproteobacteria, Fusobacteriia and Negativicutes **(Figure 2).** Class Bacteroidia represented 28% of the core microbiome, followed by Class Bacilli 19% while the least was class Flavobacteriia that represented only 0.2% of the core microbiome.

**Figure 2:**
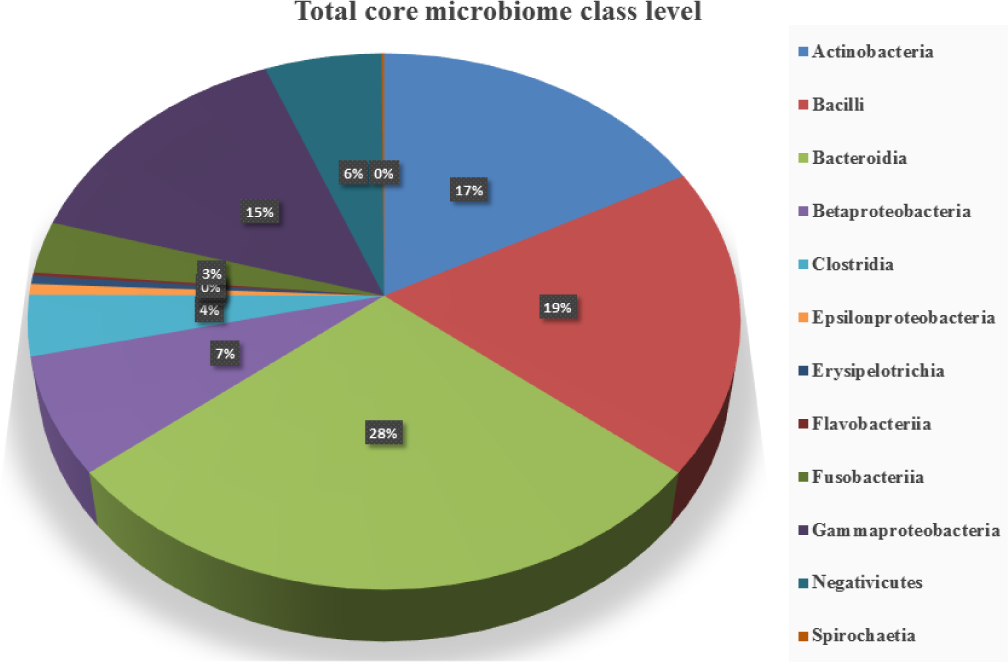
The core microbiome (class level) for the total studied population (All the 44 samples).

Similarly, for the normoglycemic group, the number of observed class that represents the core taxa (95% of reads) was also 11 classes. Class Bacteroidia represented 31% of the core microbiome, followed by class Bacilli 21.1% while the least was class Flavobacteriia that represented only 0.25% of the core microbiome. For the impaired glucose tolerance group, the same number and identities of classes, that represents the core taxa (95% of reads), also observed (11 classes). Once more, class Bacteroidia represented the highest percentage (26%) of the core microbiome. However, Gammaproteobacteria represented the second dominant taxa in impaired glucose tolerance group (21%) followed by Actinobacteria (17%), then class Bacilli (16.4%). Class Erysipelotrichia was the least component of the IGT core microbiome (0.22%). For the diabetic group, the number of observed class that represents the core taxa (95% of reads) was also 11 classes. Namely, Actinobacteria, Bacilli, Bacteroidia, Betaproteobacteria, Clostridia, Epsilonproteobacteria, Erysipelotrichia, Fusobacteriia, Gammaproteobacteria, Negativicutes, and Spirochaetia. In this group we observed the appearance of class Spirochaetia for the first time in the core microbiome and the disappearance of class Flavobacteriia. Again, class Bacteroidia represented the highest percentage (28.5%) of the core microbiome, followed by class Bacilli (19.4%). class Spirochaetia (0.4%) the least component of the diabetic core microbiome **Figure 3**.

**Figure 3:**
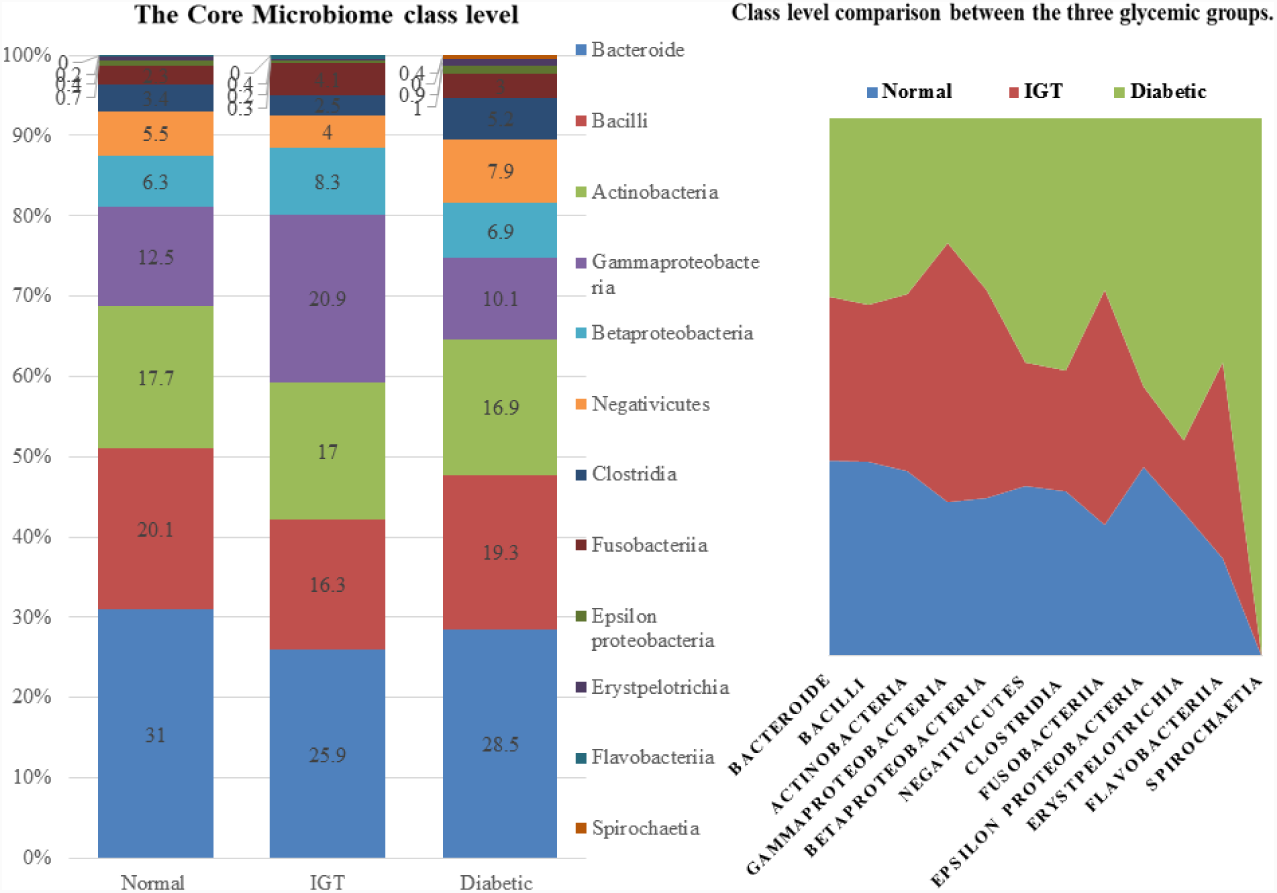
class level relative richness comparison among the three glycemic groups.

### The core bacteriome family level

For the overall sample population (44 individuals), the number of observed families that represents the core taxa (95% of reads) was 22 **(Figure 4)**. Family Prevotellaceae presented the highest percentage of the core microbiome (25.26%) while family Clostridiales XI. Incertae Sedis presented the lowermost percentage (0.17%).

**Figure 4:**
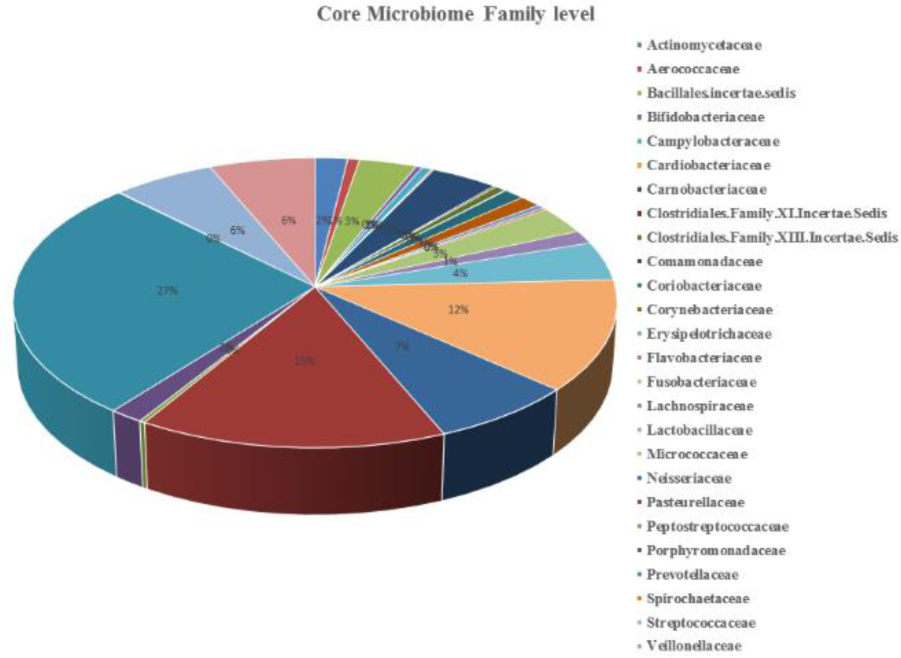
The core microbiome (family level) for the total studied population (All the 44 samples).

For the normoglycemic group, the number of observed families that represents the core taxa (95% of reads) was 24. Family Prevotellaceae presented the highest percentage of the core microbiome (29.41%) while family Clostridiales XI. Incertae Sedis presented the lowermost percentage (0.14%). For the impaired glucose tolerance group, the number of observed families that represents the core taxa (95% of reads) was 23. Family Prevotellaceae presented the highest percentage of the core microbiome (24.2%) while family Peptostreptococcaceae presented the lowermost percentage (0.16%). For the diabetic group, the number of observed families that represents the core taxa (95% of reads) was 22. Family Prevotellaceae presented the highest percentage of the core microbiome (27%) while family Peptostreptococcaceae presented the lowermost percentage (0.23%) **(Figure 5 and 6)**.

**Figure 5:**
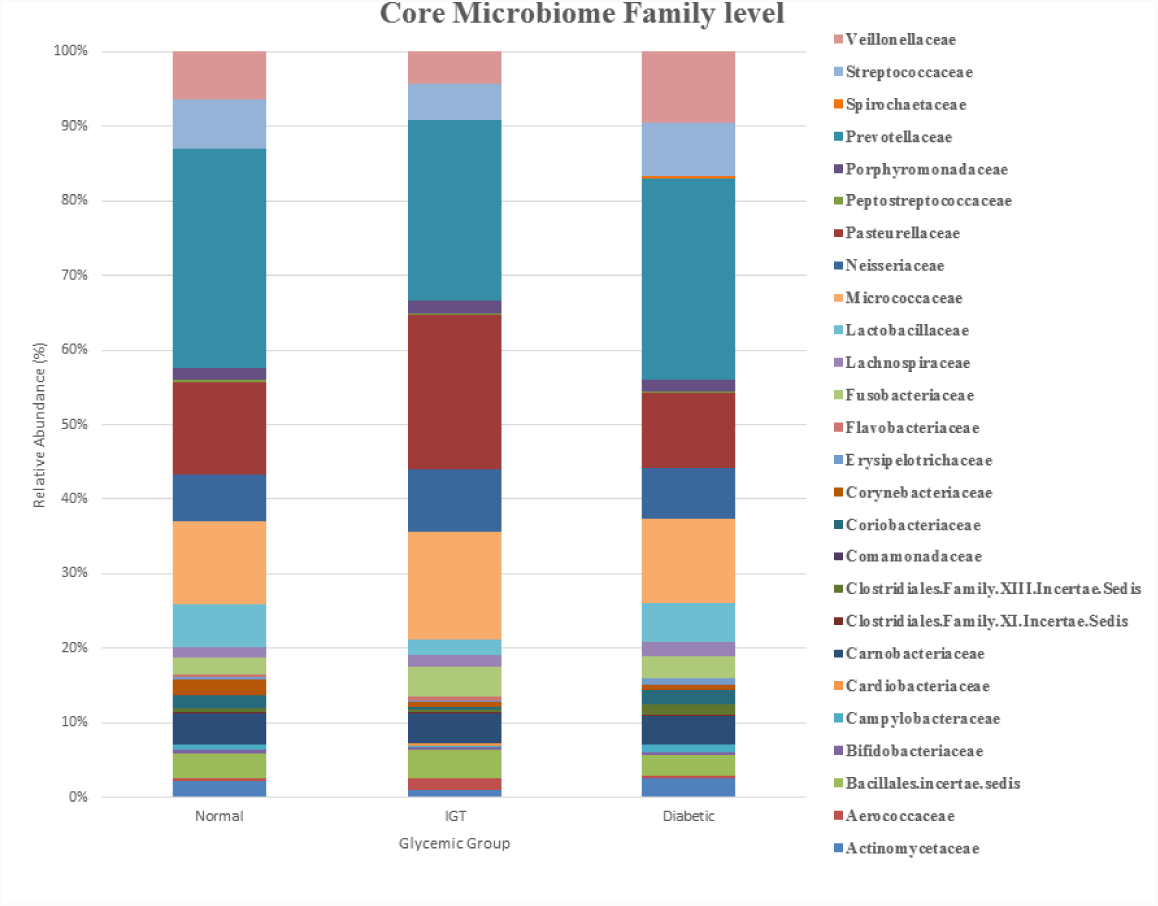
The core microbiome (family level) for the three groups.

**Figure 6:**
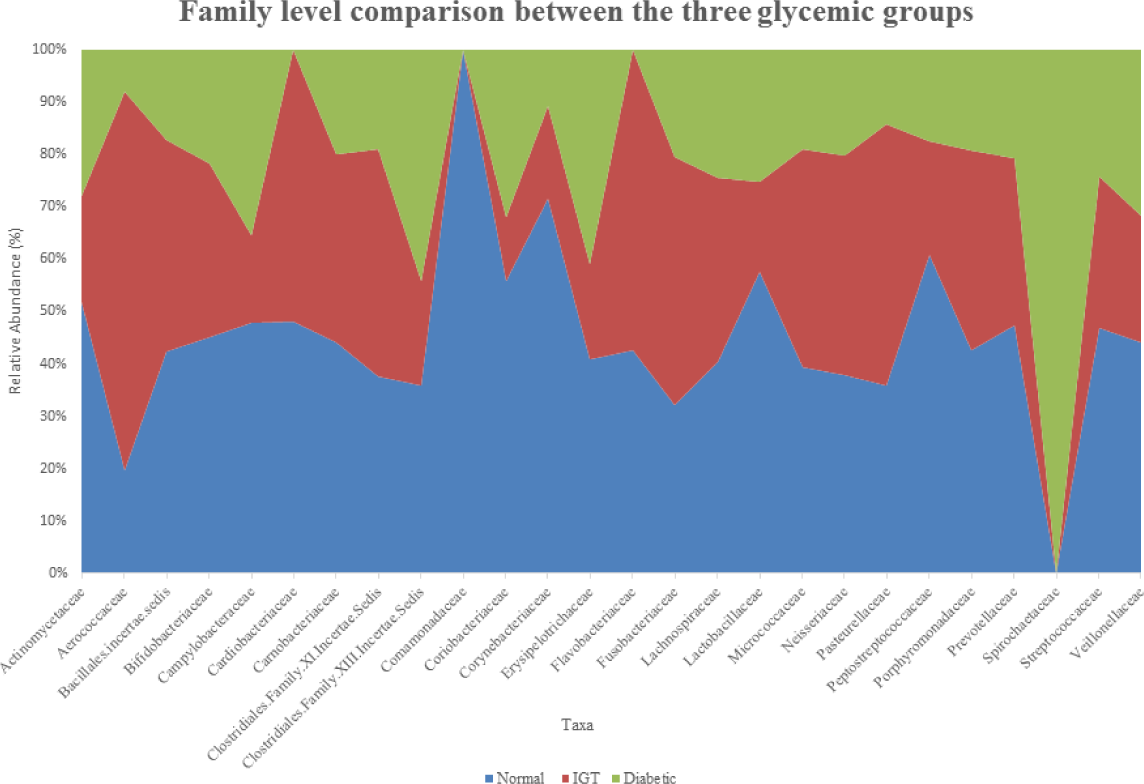
Family level comparison between the three glycemic groups.

### The core bacteriome genera level

For the overall sample population (44 individuals), the number of observed genera that represents the core taxa (95% of reads) was 29 **(Figure 7)**. Genus Prevotella presented the highest percentage of the core microbiome (26.25%) while genus Selenomonas presented the lowermost percentage (0.14%).

**Figure 7:**
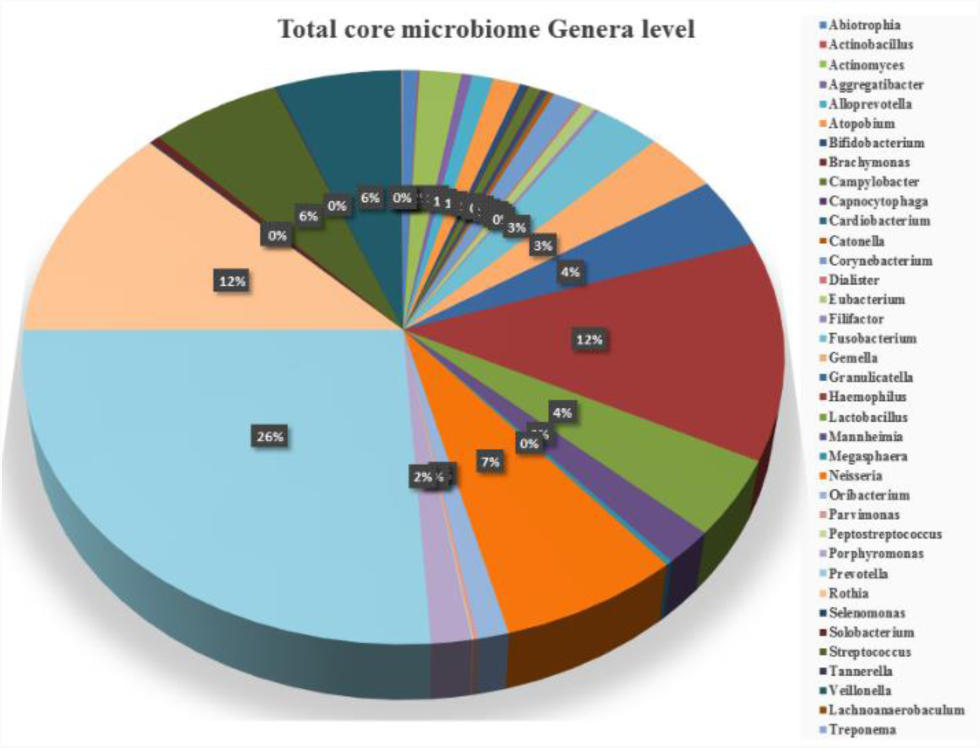
The core microbiome (genera level) for the total studied population (All the 44 samples).

For the normoglycemic group, the number of observed genera that represents the core taxa (95% of reads) was 34. Genus Prevotella presented the highest percentage of the core microbiome (28.29%), followed by genus Streptococcus (23.35%), whereas genus Peptostreptococcus presented the lowest percentage (0.12%). For the IGT group, the number of observed genera that represents the core taxa (95% of reads) was 27. Genus Prevotella presented the highest percentage of the core microbiome (23.41%), followed by genus Haemophilus (17.75%), whereas genus Filifactor presented the lowest percentage (0.16%). For the diabetic group, the number of observed genera that represents the core taxa (95% of reads) was 29. Genus Prevotella presented the highest percentage of the core microbiome (25.83%), followed by genus Rothia (11.26%), whereas genus Lachnoanaerobaculum presented the lowest percentage (0.16%).

**Figure 8:**
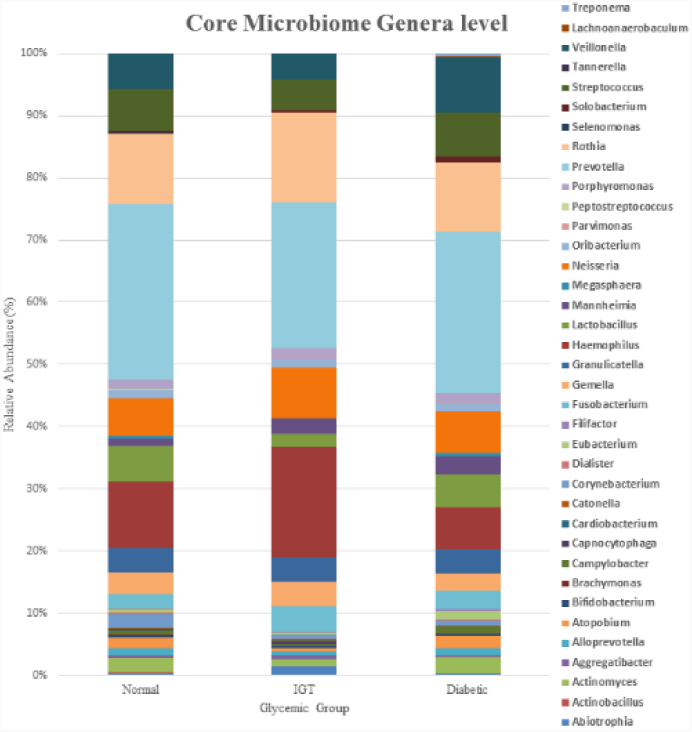
The core microbiome (Genera level) for the three groups.

**Figure 9:**
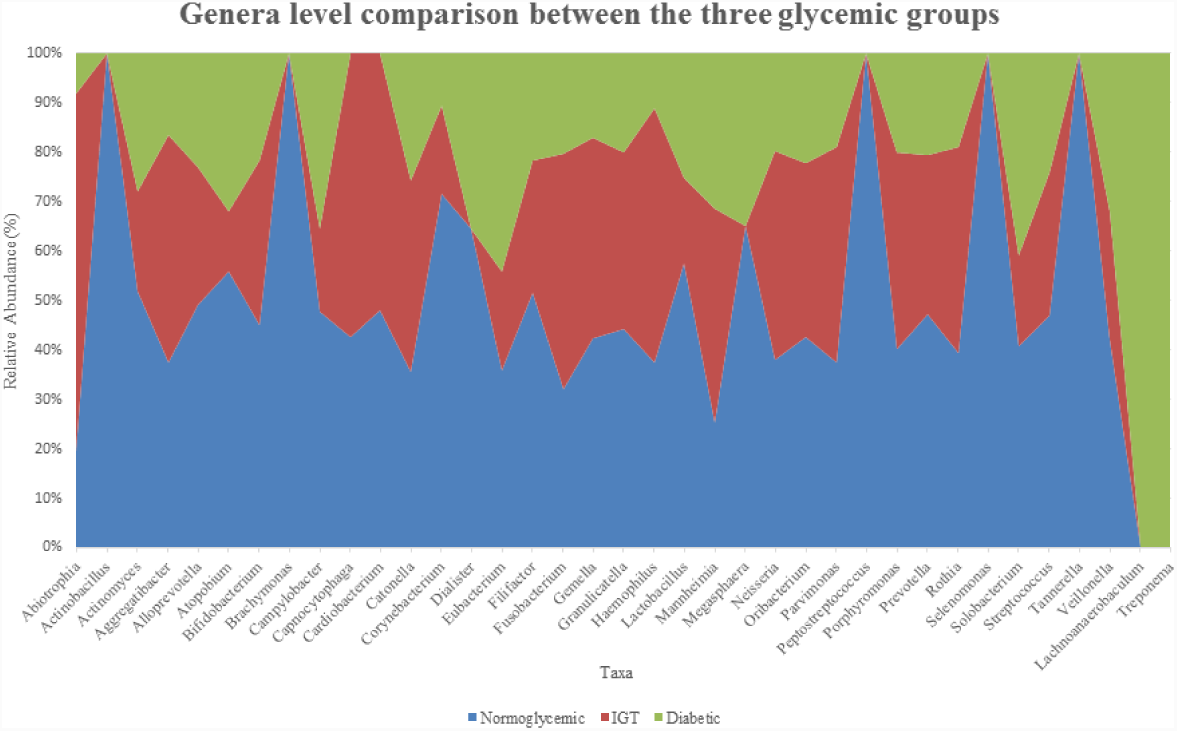
Genera level comparison between the three glycemic groups.

### The core bacteriome species level

For the overall sample population (44 individuals), the number of observed species that represents the core taxa (95% of reads) was 77 **(Figure 10)**. Species *Prevotella melaninogenica* presented the highest percentage of the core microbiome (11.68%), followed by *Rothia mucilaginosa* (11.32%), then *Haemophilus parainfluenzae* (11.1%) while species *Selenomonas noxia* presented the lowermost percentage (0.14%).

**Figure 10:**
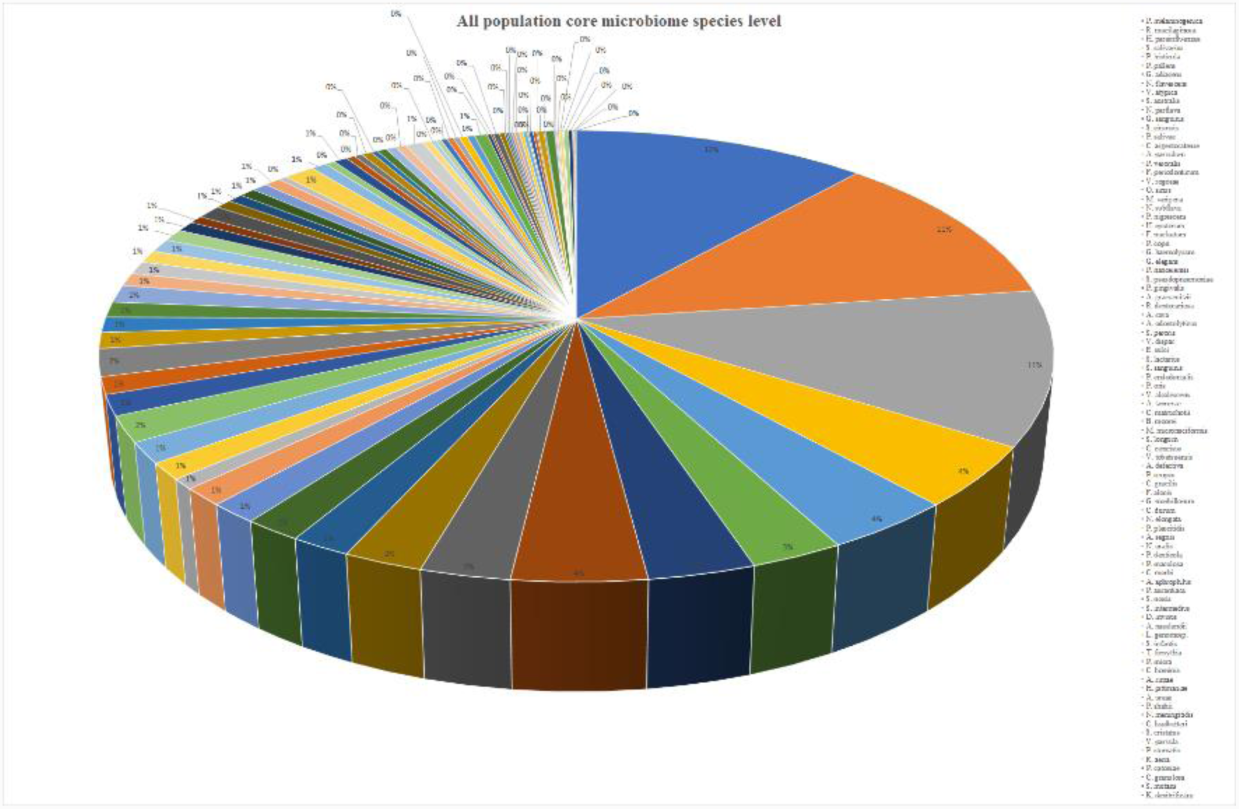
The core microbiome (species level) for the total studied population (All the 44 samples).

For the normoglycemic group, the number of observed species that represents the core taxa (95% of reads) was 87. Species *Prevotella melaninogenica* presented the highest percentage of the core microbiome (13.19%), followed by *Rothia mucilaginosa* (10.34%), then *Haemophilus parainfluenzae* (9.42%) while species *Actinomyces denitrificans* presented the lowermost percentage (0.11%).

For the IGT group, the number of observed species that represents the core taxa (95% of reads) was 71. Species *Haemophilus parainfluenzae* presented the highest percentage of the core microbiome (15.83%), followed by *Rothia mucilaginosa* (13.04%), then *Prevotella melaninogenica* (10.87%) while species *Streptococcus infantis* presented the lowermost percentage (0.15%).

For the diabetic group, the number of observed species that represents the core taxa (95% of reads) was 76 Species *Rothia mucilaginosa* presented the highest percentage of the core microbiome (10.12%), followed by *Prevotella melaninogenica* (9.61%), *Haemophilus parainfluenzae* (6.12%), then *Streptococcus salivarius* (5.35%). Whereas, species *Prevotella oulorum* presented the lowermost percentage (0.12%).

**The Supplemental Figure 2** presents the taxonomic illustration of the common core microbiome taxa distribution in, overall population, normoglycemic group, impaired glucose tolerance group and diabetic group in the mentioned above taxonomic levels. In addition **(Figure 11)** reflect the quantitative comparison of the 88 species for the core microbiome amongst the three glycemic groups. Moreover, **Figure 12** presents a core microbiome (species level) comparison between the three glycemic groups.

**Figure 11:**
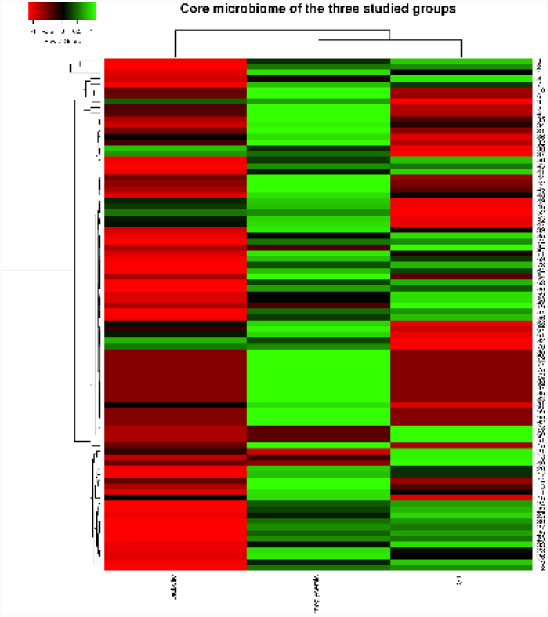
Heatmap of core species. The 88-core species are ordered by their relative abundance across the study groups. The map shows the relative abundance, normalized and log2-transformed, of the species within each group. The groups are clustered accordingly.

**Figure 12:**
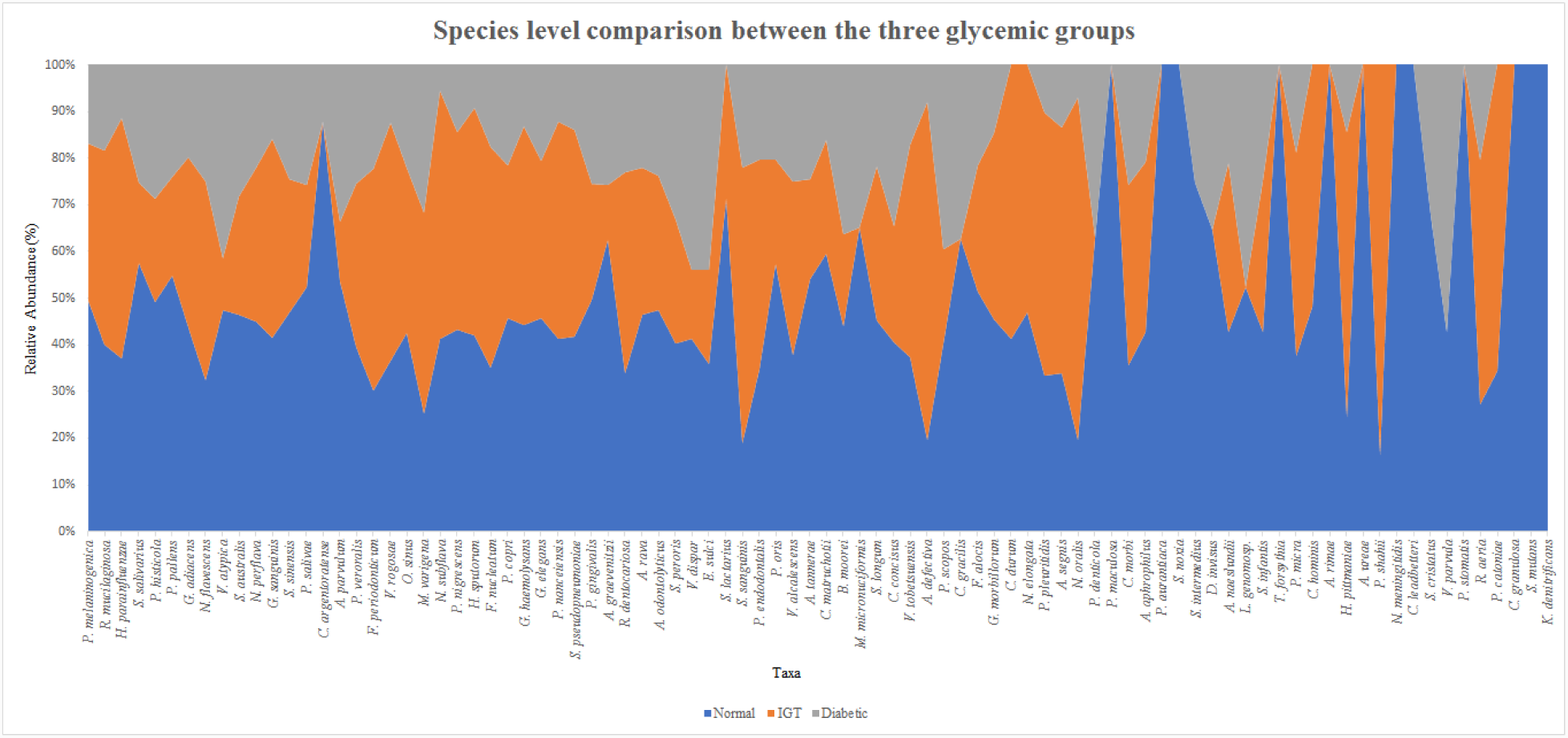
core microbiome (species level) comparison between the three glycemic groups.

### Biological diversity amongst different glycemic groups

**Table 1** presents the biological alfa diversity information and diversity indices values calculated among the three-investigated glycemic control. As we stated before the number of observed OTUs/species was the highest in the normoglycemic group (502), followed by the IGT group (412) then the diabetic group. Amongst the calculated diversity indices, the most well-known and accepted indices are Chao-1, Margalef and Fisher alpha showed that the biological diversity decrease from normoglycemic status toward the diabetic group. The Chao-1 value for the normoglycemic group is 502, IGT group is 412, then the diabetic group is 372. The Margalef index value for the normoglycemic group is 35.2, IGT group is 29.27, then diabetic group 27.49. The Fisher alpha index value for the normoglycemic group is 48.49, IGT group is 39.78, then the diabetic group is 37.71.

**Table 1:**
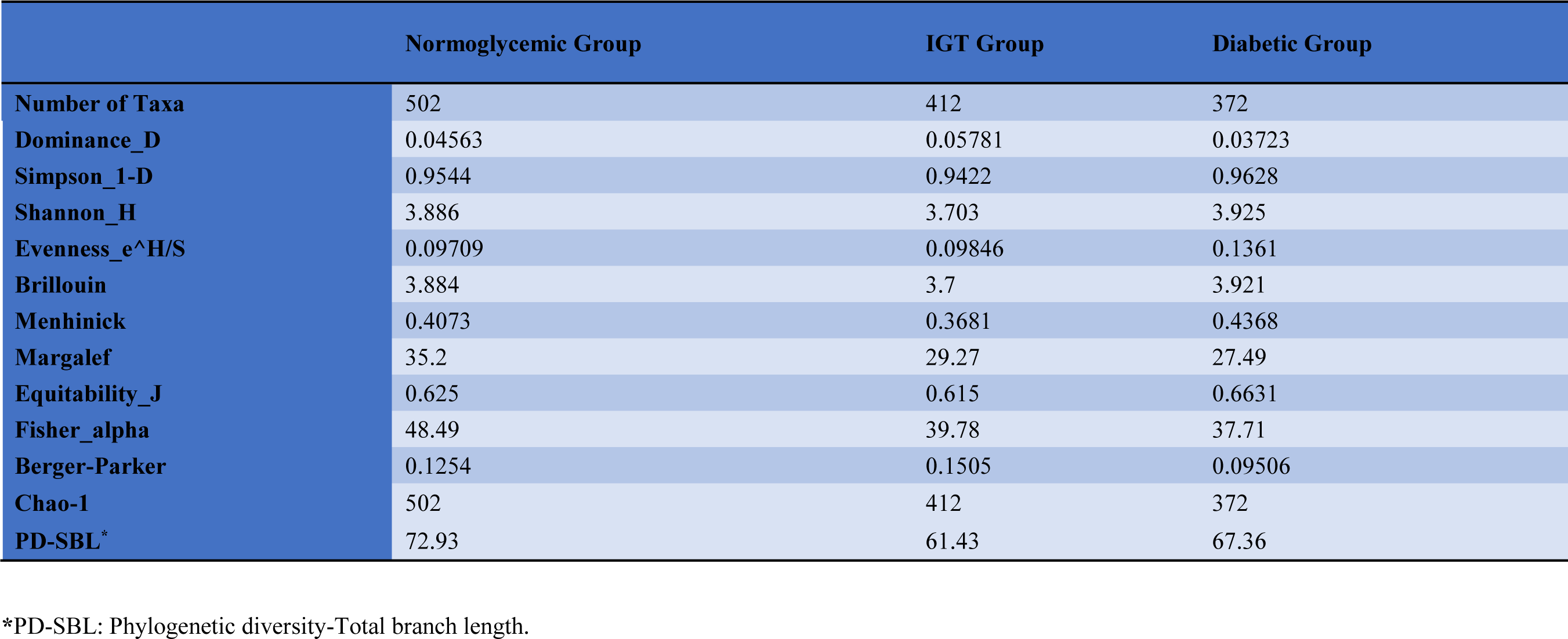
Diversity indices for the total microbiome for the three glycemic groups.

|  | Normoglycemic Group | IGT Group | Diabetic Group |
| --- | --- | --- | --- |
| **Number of Taxa** | 502 | 412 | 372 |
| **Dominance_D** | 0.04563 | 0.05781 | 0.03723 |
| **Simpson_1-D** | 0.9544 | 0.9422 | 0.9628 |
| **Shannon_H** | 3.886 | 3.703 | 3.925 |
| **Evenness_e^H/S** | 0.09709 | 0.09846 | 0.1361 |
| **Brillouin** | 3.884 | 3.7 | 3.921 |
| **Menhinick** | 0.4073 | 0.3681 | 0.4368 |
| **Margalef** | 35.2 | 29.27 | 27.49 |
| **Equitability_J** | 0.625 | 0.615 | 0.6631 |
| **Fisher_alpha** | 48.49 | 39.78 | 37.71 |
| **Berger-Parker** | 0.1254 | 0.1505 | 0.09506 |
| **Chao-1** | 502 | 412 | 372 |
| **PD-SBL\*** | 72.93 | 61.43 | 67.36 |

\*PD-SBL: Phylogenetic diversity-Total branch length.

Moreover, the beta diversity values among the three groups were as follows, Whittaker index value was 0.28538, Harrison index value was 0.14269, Cody index value was 141, Routledge index value was 0.074622, Wilson-Shmida index values was 0.32893, Mourelle index values was 0.16446, Harrison 2 index value was 0.048805 and Williams index value was 0.088929. Moreover, the pairwise Cody index value between Normoglycemic and IGT groups was 68, between Normoglycemic and diabetic groups was 99, and between IGT and diabetic was 73. We also calculated the pairwise Diversity T-tests (Shannon index) between the three tested groups. First between Normal and IGT groups. The Shannon H index values were 3.8865 and 3.7029, respectively with a t-test value of 93.427 and df: 2.6587E06 and a p-value of 0. Secondly, between Normal and diabetic groups. The Shannon H index values were 3.8865 and 3.9248, respectively with a t-test value of −17.815 and df: 1.5738E06 and a p-value of 0. On the contrary, table 1 and figure 13 showed that the lowest evenness index (Evenness_e^H/S) value was observed with the normoglycemic group (0.09709), followed by IGT group (0.09846) then the Diabetic group (0.1361).

**Figure 13:**
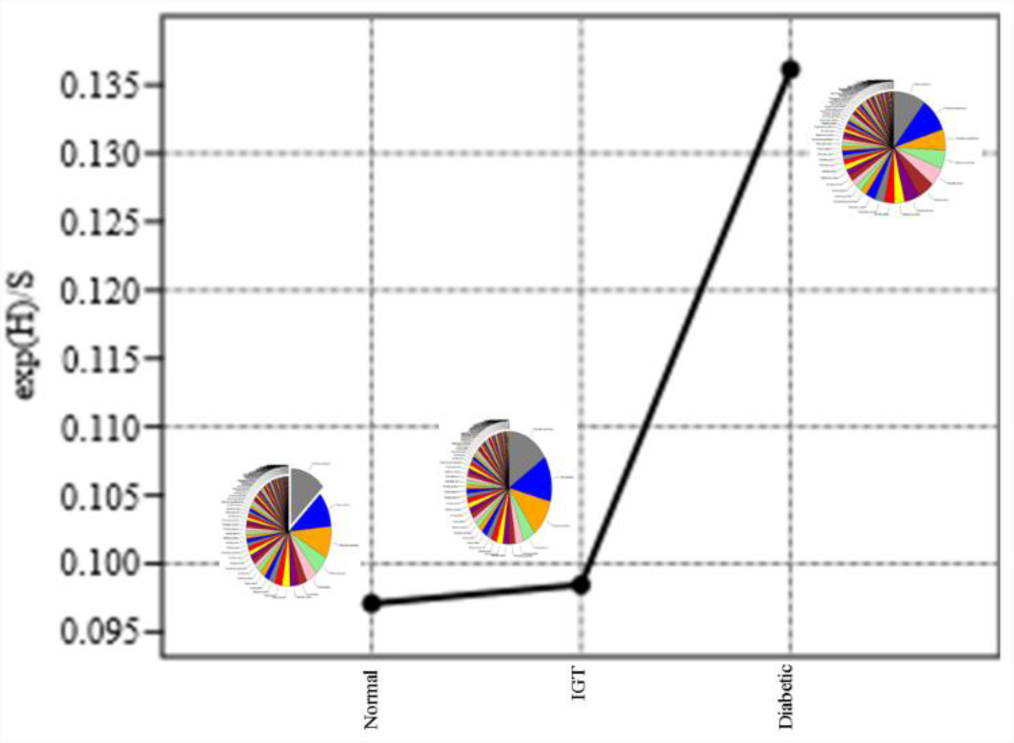
evenness index (Evenness_e^H/S) value for the three glycemic groups.

### Biological diversity within different glycemic groups

Table 2 presents the biological diversity information and diversity indices for individuals total microbiome of the normoglycemic group. The observed number of OTUs/species within the normoglycemic group ranged between 317-440. Whereas the Chao-1 index values within this group ranged from 317 to 440. The Margalef index values within this group ranged from 27.98 to 38.88. While the Fisher alpha values within this group ranged from 41.92 to 61.3. In addition, Table 3 presents the biological diversity information and diversity indices for individuals total microbiome of the IGT group. The observed number of OTUs/species within the normoglycemic group ranged between 280 and 359. Whereas the Chao-1 index values within this group ranged from 280-359. The Margalef index values within this group ranged from 23.76-30.51. While the Fisher alpha values within this group ranged from 34.08-45.32.

**Table 2:** Diversity indices for the total microbiome of the normoglycemic group.

|  | N1 | N2 | N3 | N4 | N5 | N6 | N7 | N8 | N9 | N10 | N11 | N12 | N13 | N14 | N15 | N16 | N17 | N18 | N19 |
| --- | --- | --- | --- | --- | --- | --- | --- | --- | --- | --- | --- | --- | --- | --- | --- | --- | --- | --- | --- |
| Number of OTUs | 405 | 317 | 377 | 374 | 390 | 317 | 352 | 370 | 390 | 339 | 354 | 328 | 378 | 440 | 325 | 350 | 344 | 361 | 361 |
| Individuals | 78853 | 80404 | 79615 | 79688 | 79741 | 80436 | 79696 | 79032 | 78831 | 80103 | 80103 | 80261 | 79671 | 80009 | 80480 | 80177 | 80363 | 80041 | 79148 |
| Dominance_D | 0.0567 | 0.09204 | 0.04905 | 0.0712 | 0.06713 | 0.1421 | 0.05888 | 0.05326 | 0.05079 | 0.06644 | 0.05731 | 0.1118 | 0.06402 | 0.04983 | 0.08974 | 0.05824 | 0.1046 | 0.0358 | 0.07406 |
| Simpson_1-D | 0.9433 | 0.908 | 0.9509 | 0.9288 | 0.9329 | 0.8579 | 0.9411 | 0.9467 | 0.9492 | 0.9336 | 0.9427 | 0.8882 | 0.936 | 0.9502 | 0.9103 | 0.9418 | 0.8954 | 0.9642 | 0.9259 |
| Shannon_H | 3.866 | 3.033 | 3.715 | 3.569 | 3.404 | 2.577 | 3.49 | 3.819 | 3.846 | 3.339 | 3.557 | 3.032 | 3.629 | 3.616 | 3.077 | 3.491 | 3.062 | 3.856 | 3.347 |
| Evenness_e^H/S | 0.1179 | 0.06551 | 0.1089 | 0.09484 | 0.07717 | 0.04151 | 0.09314 | 0.1231 | 0.1201 | 0.08317 | 0.09903 | 0.06323 | 0.09971 | 0.08449 | 0.06676 | 0.09375 | 0.06215 | 0.1309 | 0.07871 |
| Brillouin | 3.822 | 3 | 3.673 | 3.528 | 3.363 | 2.543 | 3.453 | 3.778 | 3.804 | 3.303 | 3.519 | 2.999 | 3.587 | 3.57 | 3.043 | 3.455 | 3.028 | 3.818 | 3.326 |
| Menhinick | 1.44 | 1.117 | 1.334 | 1.323 | 1.379 | 1.116 | 1.245 | 1.314 | 1.387 | 1.196 | 1.249 | 1.157 | 1.337 | 1.553 | 1.144 | 1.235 | 1.212 | 1.274 | 1.283 |
| Margalef | 35.83 | 27.98 | 33.32 | 33.05 | 34.47 | 27.98 | 31.1 | 32.72 | 34.5 | 29.94 | 31.26 | 28.96 | 33.41 | 38.88 | 28.68 | 30.91 | 30.37 | 31.89 | 31.92 |
| Equitability_J | 0.6439 | 0.5267 | 0.6262 | 0.6024 | 0.5706 | 0.4475 | 0.5952 | 0.6457 | 0.6447 | 0.5731 | 0.606 | 0.5234 | 0.6115 | 0.594 | 0.532 | 0.5959 | 0.5243 | 0.6548 | 0.5683 |
| Fisher_alpha | 55.81 | 41.92 | 51.29 | 50.81 | 53.33 | 41.92 | 47.37 | 50.24 | 53.42 | 45.32 | 47.64 | 43.61 | 51.44 | 61.3 | 43.14 | 47.01 | 46.07 | 48.74 | 48.84 |
| Berger-Parker | 0.1713 | 0.1967 | 0.121 | 0.2045 | 0.1519 | 0.229 | 0.1252 | 0.1581 | 0.1551 | 0.1368 | 0.1589 | 0.2741 | 0.1603 | 0.1183 | 0.225 | 0.1449 | 0.2317 | 0.1058 | 0.1725 |
| Chao-1 | 405 | 317 | 377 | 374 | 390 | 317 | 352 | 370 | 390 | 339 | 354 | 328 | 378 | 440 | 325 | 350 | 344 | 361 | 361 |
| PD-SBL<sup>*</sup> | 54.14 | 54.89 | 56.74 | 56.11 | 62.67 | 46.91 | 53.50 | 55.84 | 55.95 | 52.23 | 52.23 | 53.84 | 54.41 | 62.89 | 52.64 | 53.63 | 55.17 | 52.44 | 50.95 |

\*Phylogenetic diversity-Total branch length.

**Table 3:** Diversity indices for the total microbiome of the Impaired glucose tolerance group.

|  | IGT1 | IGT2 | IGT3 | IGT4 | IGT5 | IGT6 | IGT7 | IGT8 | IGT9 | IGT10 |
| --- | --- | --- | --- | --- | --- | --- | --- | --- | --- | --- |
| Number of OTUs | 303 | 359 | 319 | 315 | 331 | 350 | 326 | 322 | 317 | 280 |
| Individuals | 125219 | 124698 | 125958 | 122766 | 125099 | 124609 | 125984 | 125762 | 126100 | 125947 |
| Dominance_D | 0.05302 | 0.04789 | 0.09629 | 0.06048 | 0.06958 | 0.06733 | 0.1065 | 0.06911 | 0.07404 | 0.1246 |
| Simpson_1-D | 0.947 | 0.9521 | 0.9037 | 0.9395 | 0.9304 | 0.9327 | 0.8935 | 0.9309 | 0.926 | 0.8754 |
| Shannon_H | 3.419 | 3.791 | 3.148 | 3.467 | 3.489 | 3.644 | 2.974 | 3.43 | 3.377 | 2.809 |
| Evenness_e^H/S | 0.1008 | 0.1234 | 0.07301 | 0.1017 | 0.09897 | 0.1092 | 0.06003 | 0.09585 | 0.09237 | 0.05926 |
| Brillouin | 3.4 | 3.771 | 3.13 | 3.449 | 3.471 | 3.624 | 2.957 | 3.416 | 3.359 | 2.794 |
| Menhinick | 0.8558 | 1.016 | 0.8984 | 0.8986 | 0.9354 | 0.991 | 0.918 | 0.9077 | 0.8922 | 0.7886 |
| Margalef | 25.73 | 30.51 | 27.08 | 26.8 | 28.12 | 29.75 | 27.67 | 27.34 | 26.91 | 23.76 |
| Equitability_J | 0.5985 | 0.6444 | 0.546 | 0.6027 | 0.6014 | 0.622 | 0.5139 | 0.5939 | 0.5864 | 0.4985 |
| Fisher_alpha | 37.32 | 45.32 | 39.54 | 39.12 | 41.28 | 44.03 | 40.53 | 39.98 | 39.25 | 34.08 |
| Berger-Parker | 0.1097 | 0.1324 | 0.2479 | 0.137 | 0.1605 | 0.2172 | 0.2292 | 0.1903 | 0.1896 | 0.2093 |
| Chao-1 | 303 | 359 | 319 | 315 | 331 | 350 | 326 | 322 | 317 | 280 |
| PD-SBL* | 54.22 | 57.58 | 54.32 | 46.99 | 52.84 | 56.85 | 43.98 | 52.08 | 55.62 | 47.43 |

\* Phylogenetic Diversity-Total branch length.

Furthermore, Table 4 presents the biological diversity information and diversity indices for individuals total microbiome of the diabetic group. The observed number of OTUs/species within the normoglycemic group ranged between 221-289. Whereas the Chao-1 index values within this group ranged from 221 to 289. The Margalef index values within this group ranged from 20.41-27.56. While the Fisher alpha values within this group ranged from 29.94 to 42.36. In addition, Figure 14 presents the biological diversity using Chao-1 diversity index presented in a box plot illustration. The mean value for normal subjects was 361±32, while IGT patients had a mean of 322±22 and 262±18 was the mean for the diabetic population. The difference between the 2 groups was significant with a p-value <0.001.

**Table 4:**
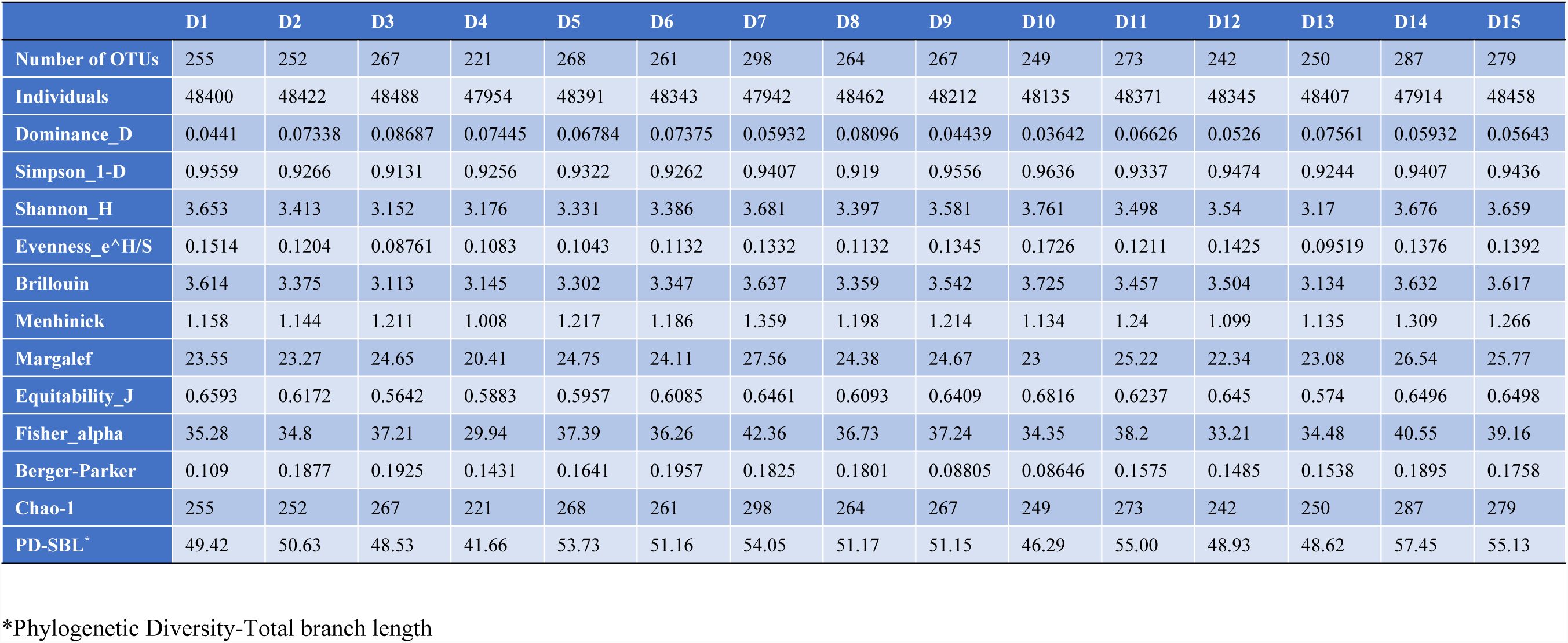
Diversity indices for the total microbiome of the diabetic group.

**Figure 14:**
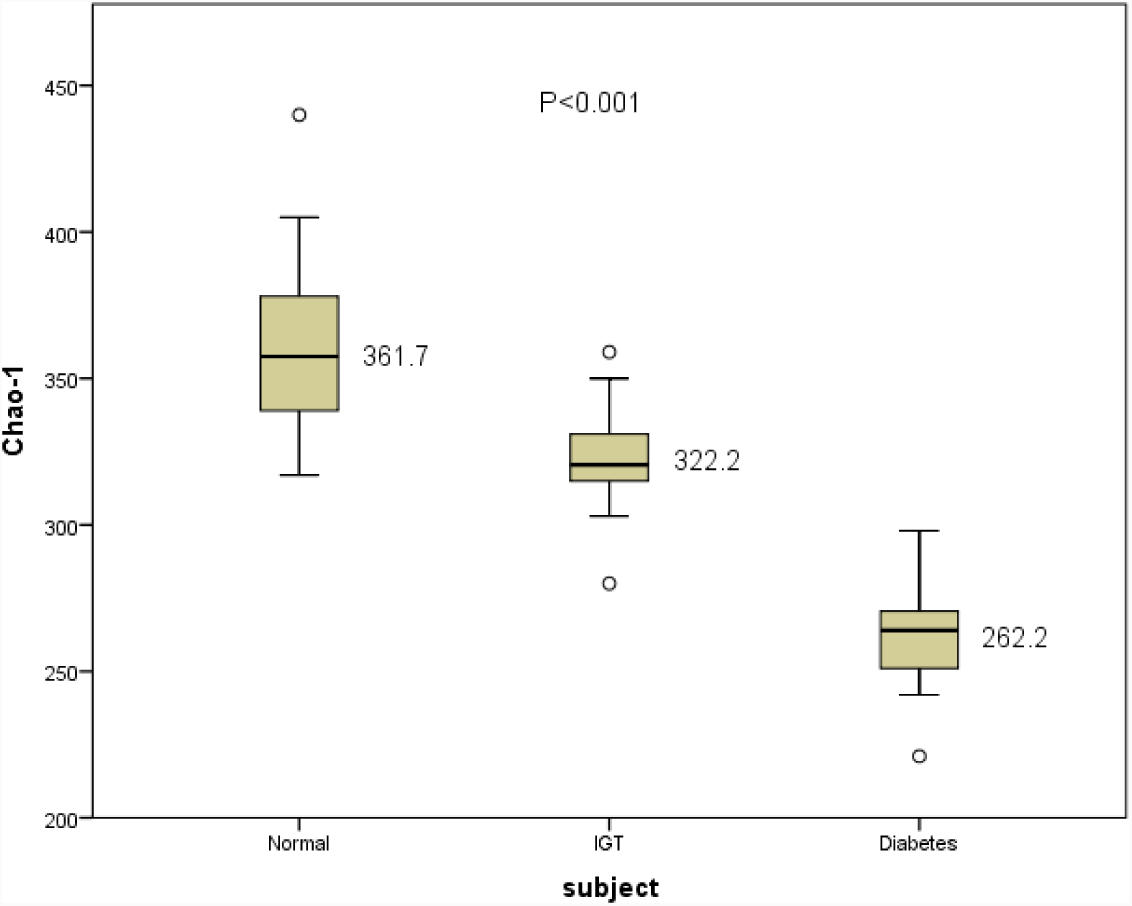
biological diversity using Chao-1 diversity index presented in a box plot depiction.

In addition, figures 15, 16 and 17 reflects the quantitative comparison of the 88 species for the core microbiome amongst the individuals of the three different glycemic groups namely, normal, IGT and diabetic respectively.

**Figure 15:**
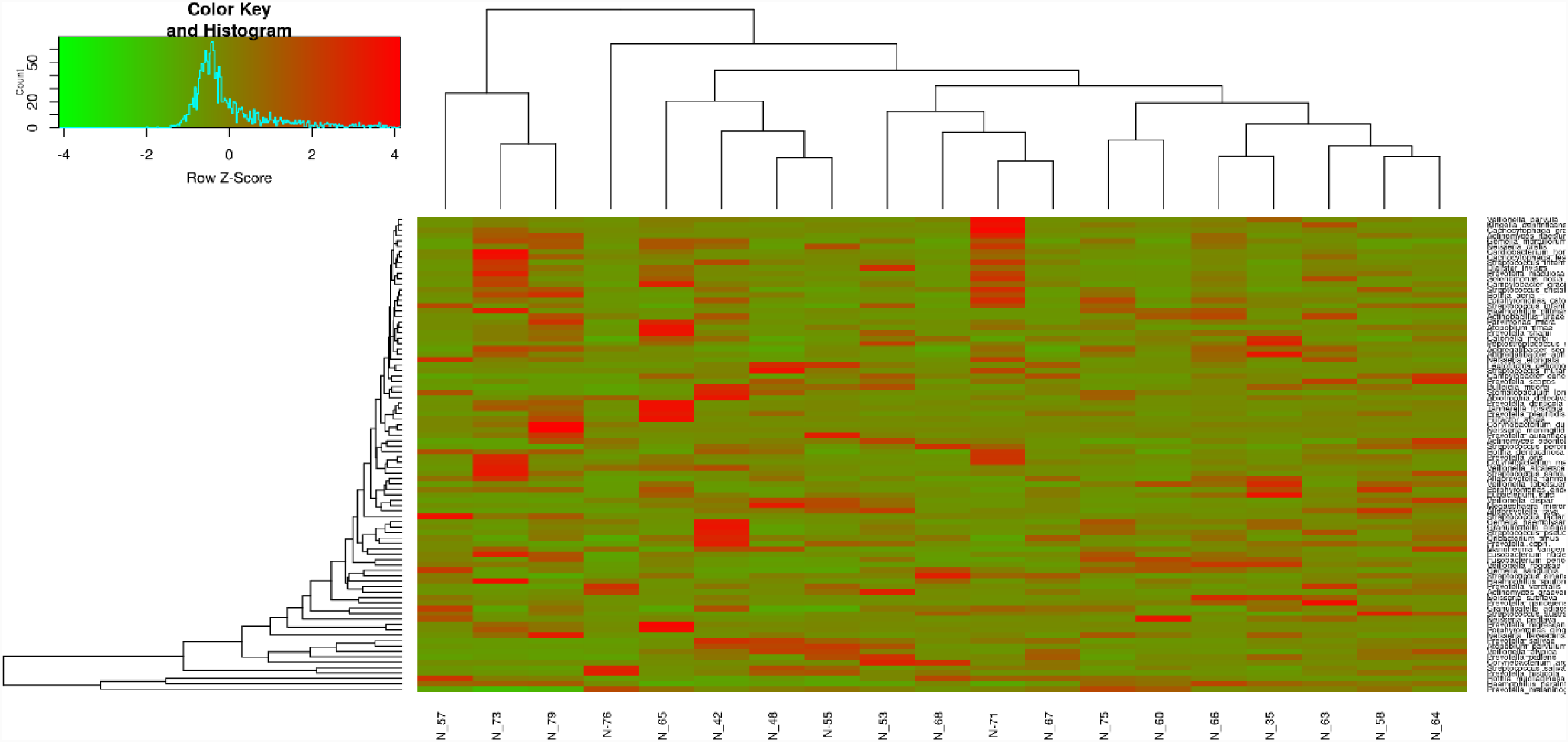
Heatmap of core species. The 88-core species are ordered by their relative abundance across the normoglycemic subjects. The map shows the relative abundance, normalized and log2-transformed, of the species within each subject. The subjects are clustered accordingly.

**Figure 16:**
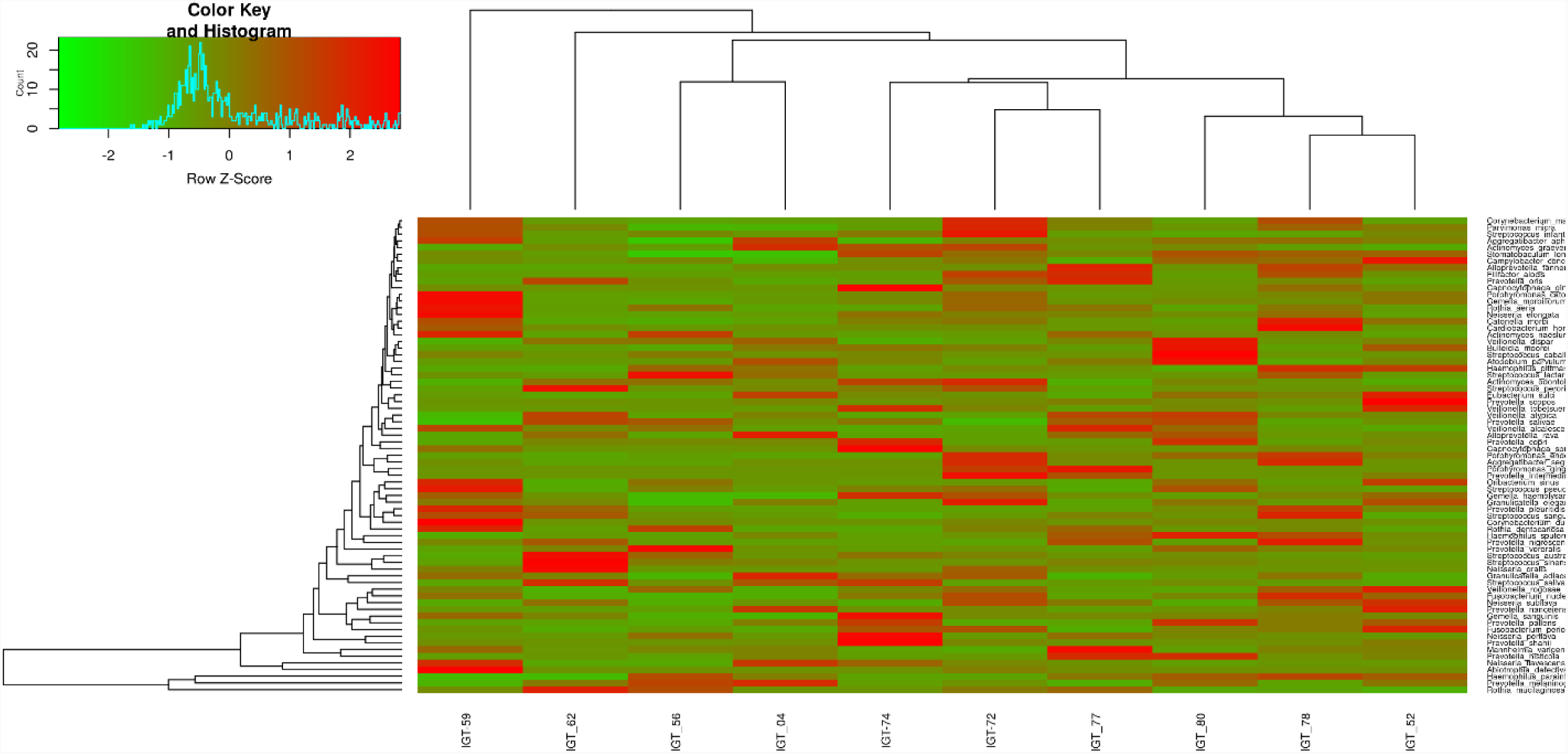
Heatmap of core species. The 88-core species are ordered by their relative abundance across the IGT subjects. The map shows the relative abundance, normalized and log2-transformed, of the species within each subject. The subjects are clustered accordingly.

**Figure 17:**
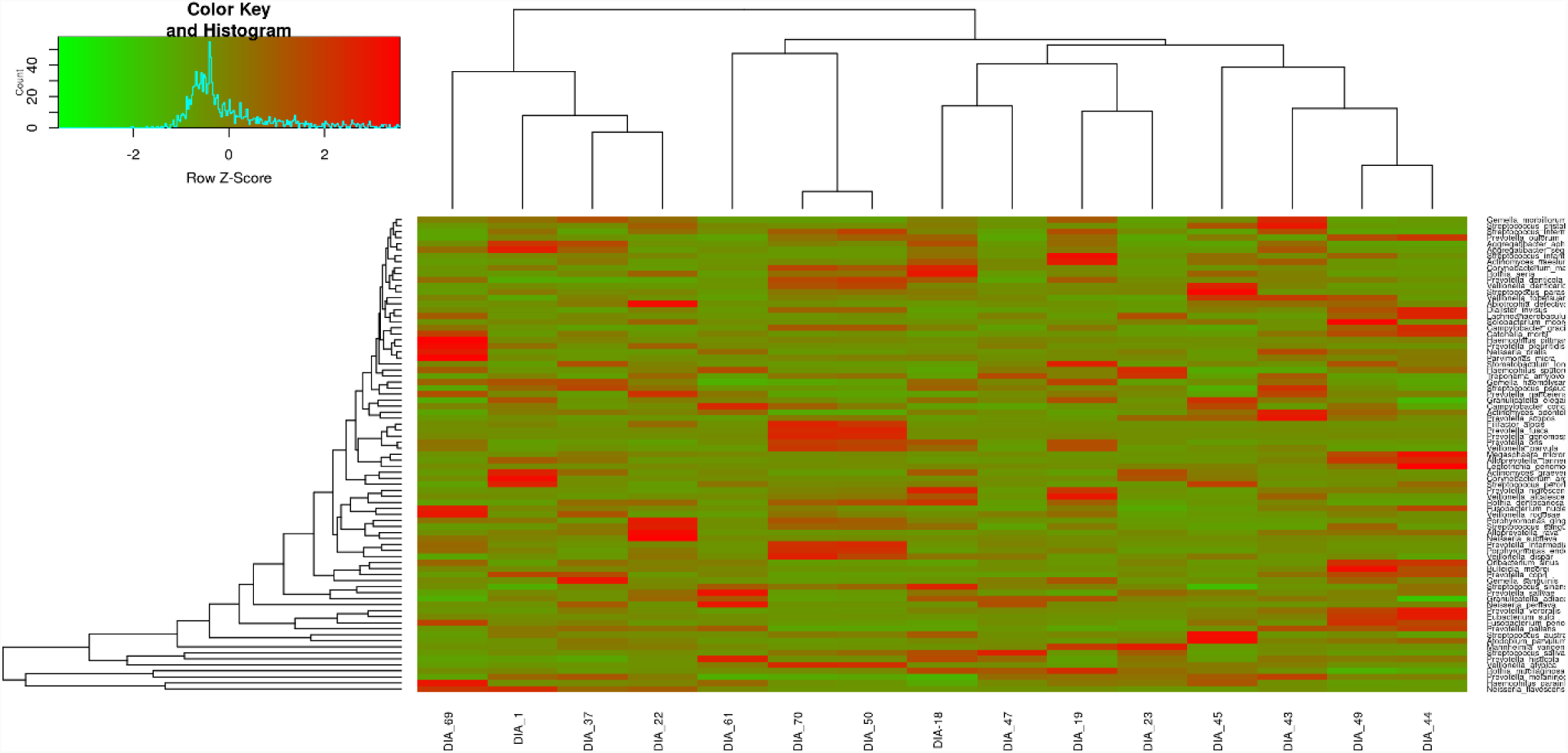
Heatmap of core species. The 87-core species are ordered by their relative abundance across the diabetic subjects. The map shows the relative abundance, normalized and log2-transformed, of the species within each subject. The subjects are clustered accordingly.

### Phylogenetic Diversity analysis

**Table 1 also** presents the phylogenetic diversity information (PD-SBL) for the three-investigated glycemic control. The normoglycemic Group exhibited the highest value of PD-SBL (72.93) **(The Supplemental Figure 3)**, followed by diabetic group (67.36) **(The Supplemental Figure 4)** then diabetic group IGT group (61.43) **(The Supplemental Figure 5). Table 2** presents the phylogenetic diversity information for individuals total microbiome of the normoglycemic group. The highest value of PD-SBL (62.88991276) was observed for individual 14 **(The Supplemental Figure 6)**, while the lowest value of PD-SBL (46.90922608) was observed for individual 6 **(The Supplemental Figure 7)**. **Table 3** presents the phylogenetic diversity information for individuals total microbiome of the IGT group. The highest value of PD-SBL (57.58284984) was observed for subject 2 **(The Supplemental Figure 8),** whereas the lowest value of PD-SBL (43.97685086) was observed for subject 7 **(The Supplemental Figure 9). Table 4** presents the phylogenetic diversity information for individuals total microbiome of the diabetic group. The highest value of PD-SBL (57.45173230) was observed for subject 14 **(The Supplemental Figure 10),** whereas the lowest value of PD-SBL (41.65727144) was observed for subject 4 **(The Supplemental Figure 11).**

## Discussion

In this study we selected diabetic and IGT patients to represent abnormal glucose metabolism conditions. We utilized a very strict criterion to select individual participating in this investigation. The normal control subjects were selected from the patient’s immediate family members to account for host genetic background and environmental effects which are very important factors for host-associated microbial communities. It was found that oral microbiome similarity increased with shared host genotype, irrespective of dental state. In addition, it was found that environmental factors have also a great influence on the oral microbiome, however, many taxa are inherited from the parents. On the other hand, possibly cariogenic bacterial taxa are likely not controlled by genetic factors [26]. In our study we selected subjects and diabetic patients that are either free of any periodontitis or having mild periodontitis to diminish the effect of oral health on the oral microbiome structure and attempt to make any observed changes in the oral flora is due to the glycemic status only as much as possible. Clear changes in the oral flora were found in diabetic patients with both aggressive or chronic periodontitis [11] [27] [28]. In this study, there is a clear reduction in the number of species (OTUs) observed in both impaired glucose tolerance and in the diabetic group when compared with the normoglycemic subjects. This could be explained by three different mechanisms. Firstly, the presence of higher glucose concentration in the saliva as a change the microbiome environment may enhance certain bacterial species on the expense of others that cause the reduced microbiome numbers observed in diabetic and pre-diabetic [29] [30]. Secondly, the presence of mouth dehydration that is usually associated with the diabetic condition can cause the observed microbial diversity reduction [31]. The third explanation is related to acidification of the oral environment because of hyperglycemia leading to generalized perturbation in the oral microbiome [32].

In general, we noticed that more than 50% of the total observed species were shared by the three clinical groups, while 20% were shared by any two of those clinical conditions and the remaining 30% were distributed in a single group **(Figure 1)** and (**Supplementary Table 2**). However, we observed 86 OTUs only in the normoglycemic group, 15 OTUs only in the IGT group and 26 OTUs only in the diabetic group. In addition, we observed 78 shared OTUs between the normal and IGT groups, 27 between Normal and diabetic groups and 8 between IGT and diabetic groups. These shared OTUs considered transitional taxa. We propose that these differential and transitional species associated with each clinical state could have a clinical value and require a closer screening. Thus, we investigated the number and types of bacterial species for being non-pathogenic, pathogenic or beneficial or probiotic bacteria. The non-pathogenic taxa are those bacteria that were not oral tissue infection or caries, while pathogenic species are those that were found to be associated with any oral infections such as periodontitis or related to dental caries. Whereas the probiotic microorganisms are these providing oral or general health benefits.

First, for the 86 OTUs exclusive to the normoglycemic group, 63 (73.3%) were not oral pathogens, 15 (17.4%) were oral/dental pathogens and 8 (9.3%) were beneficial of probiotic bacteria (**Supplementary Table 3**). For example, *Actinomyces radicidentis* was isolated in pure culture from infected root canals of teeth [33]. Moreover, *Mogibacterium neglectum* recovered from tongue plaque and necrotic dental pulp [34]. On the other hand, *Lactobacillus paracasei*, which has been established to suppress the growth of many cariogenic and pathogenic microbes such as *Streptococcus mutans*, was only observed in the normoglycemic group [35]. Moreover, we observed the presence of *Lactobacillus oris* only in the normoglycemic group. *Lactobacillus oris* is a candidate probiotic for the prevention of oral diseases [36]. One more beneficial bacterial species we observed in only in this group was *Lactobacillus crispatus. L. crispatus* is considered as a new probiotic species with probable oral health benefits with high antibacterial activity against periodontal bacteria [37].

Secondly, for the 15 OTUs exclusive to the IGT group, 7 (46.7 %) were not oral pathogens, 5 (33.3 %) were oral/dental pathogens and 3 (20%) were beneficial of probiotic bacteria. The oral/dental pathogens exclusive for IGT group were *Enterococcus gallinarum*, *Streptococcus genomosp*, *Capnocytophaga sp. Strain F0488*, *Treponema pectinovorum* and *Moraxella caprae* (**Supplementary Table 4**). *Enterococcus gallinarum* is indicated in spontaneous bacterial peritonitis [38,39]. In addition, *Streptococcus genomosp*, *Capnocytophaga sp.* Strain F0488 were indicated in caries and opportunistic oral infection, respectively [40,41]. In addition, *Treponema pectinovorum* was associated with endodontic treatment failure and periapical dental lesions [42]. However, two bacterial species with anti-inflammatory were observed in this group namely, *Weissella paramesenteroides* and *Bifidobacterium breve.* In addition, *W. paramesenteroides* produce bacteriocins against numerous pathogens [43] [44].

Third, For the 26 OTUs exclusive to the diabetic group, 16 (61.54 %) were not oral pathogens, 10 (38.46 %) were oral/dental pathogens and no (0%) beneficial probiotic bacteria were observed (**Supplementary Table 5**). Among the observed oral pathogens in the diabetic group, *Treponema pallidum*, which is highly associated with oral cavity infections in immunodeficiency virus-infected individuals and is also the oral syphilis-causing pathogen [45] [46]. *Treponema sp. Clone AF026* isolated from human periodontal pockets of several subjects with a range of periodontal conditions [47]. In addition, *Lactobacillus ultunensis* is bacterial species implicated in advanced dental caries [48] [49]. Interestingly, we observed the presence of the non-oral pathogen *Peptoniphilus asaccharolyticus* among the diabetic group. However, *P. asaccharolyticus* is associated with septic arthritis and osteomyelitis in a woman with osteoarthritis and diabetes mellitus and previously isolated from Diabetic Foot Infections [50] [51].

Normoglycemic and IGT groups shared 78 bacterial species OTUs. Among these 78-transitional species (Between phases) consisted of 10 positive oral pathogens (12.82%), 12 potential oral pathogens (15.38%), 56 non-oral pathogens (71.79%) and no beneficial probiotic bacteria were observed (**Supplementary Table 6**). Among the oral pathogens in normoglycemic/IGT transitional species (OTUs), *Actinomyces israelii* was observed. *A. israelii* is the causal agent of actinomycosis and oral periapical lesion and oral-cervicofacial (“lumpy jaw”) infection [52] [53]. Moreover, *Arcanobacterium haemolyticum*, the causal agent of Lemierre syndrome that includes oropharyngeal infection was also observed [54]. In addition, *Brevundimonas diminuta*, the causal agent of refractory periodontitis and advanced Noma lesions was also observed [55] [56].

IGT and Diabetic group shared 8 bacterial species OTUs. Among these 8-transitional species (Between phases) consisted of four positive oral pathogens (50%), one potential oral pathogen (12.5%), three non-oral pathogens (37.5%) and no beneficial probiotic bacteria were observed (**Supplementary Table 7**). Among the oral pathogens in IGT/diabetic transitional species (OTUs), *Staphylococcus warneri*, that has role apical periodontitis lesions of obturated teeth and persistent root canal infections [57] [58]. In addition, *Leptothrix_sp_Clone_AV011a*, the causal agent of advanced Noma lesions and dentine caries was also observed [56] [59]. The potential oral pathogens observed, *Streptococcus_downei*, is associated with dental plaque in humans [60]. Even though we observed three non-oral pathogens among this collection, but it can cause infections elsewhere. For example, *Haemophilus paraphrohaemolyticus* is the causal agent of liver abscess [61].

This marked increase in the pathogenic content of the hyperglycemic microbiome could be explained by the decreased immunity associated with this condition and/or the acidification of saliva as a result of the presence of glucose since those two factors are known to affect the growth of microorganisms in the oral cavity [62]. Among type 2 diabetic patients, the microbiome distribution was totally different from normal subjects or IGT patients. This group of patients had the highest rate of pathogenic microbiome at 38.5% and more non-pathogenic bacteria than IGT patients but less than normal subjects. There were no probiotic microorganisms isolated from the oral cavity of those patients. The increase in pathogenic oral microbiome observed in this group may share the same factors explained in the IGT group, although the factors will be more in this group namely: lower immunity and higher oral cavity acidity. The reasons behind the total absence of probiotic microorganisms could be the increased pathogenic microbiome which could be associated with the presence of toxic materials that would inhibit probiotic bacteria. Other factors that are related to hyperglycemia may have had contributed to this change which needs further studies. In addition, a future pathogenomics study will be required to confirm our observed results on a metagenomic level.

## Conclusion

In the present study, we observed clear reduction in the biological and phylogenetic diversity in the diabetic and pre-diabetic oral microbiome in comparison with the normoglycemic oral microbiome. However, this reduction was associated with an increase in the pathogenic content of the hyperglycemic microbiomes. A future pathogenomics, i.e. pathogenicity and virulence factors determinant, study will be essential to confirm our observed results on a metagenomic level.

## Supplemental Figures

**Supplementary Figure 1.** Rarefaction curves for the number of observed operational taxonomic units (OTUs) per sample in Normoglycemic group (a), Impaired glucose tolerance group (b) and Diabetic group (c).

**Supplementary Figure 2.** Taxonomic representation of the common core microbiome taxa distribution in a) overall population b) normoglycemic group, c) impaired glucose tolerance group, d) diabetic group.

**Supplemental Figure 3:** Phylogenetic diversity of the normoglycemic Group

**Supplemental Figure 4:** Phylogenetic diversity of the diabetic group.

**Supplemental Figure 5:** Phylogenetic diversity of the IGT group.

**Supplemental Figure 6:** Phylogenetic diversity information for individual 14 from normoglycemic group with the highest value of PD-SBL (62.88991276).

**Supplemental Figure 7:** Phylogenetic diversity information for individual 6 from normoglycemic group with the lowest value of PD-SBL (46.90922608).

**Supplemental Figure 8:** Phylogenetic diversity information for individual 2 from IGT group with the highest value of PD-SBL (57.58284984).

**Supplemental Figure 9:** Phylogenetic diversity information for individual 7 from normoglycemic group with the lowest value of PD-SBL (43.97685086).

**Supplemental Figure 10:** Phylogenetic diversity information for individual 14 from diabetic group with the highest value of PD-SBL (57.45173230).

**Supplemental Figure 11:** Phylogenetic diversity information for individual 4 from normoglycemic group with the lowest value of PD-SBL (41.65727144).

## Supplementary Tables

**Supplementary table 1:** The clinical information for the diabetic patients participated in the study.

**Supplementary table 2:** Distribution of the observed oral microbiome species (OTUs) among the different studied Glycemic groups.

**Supplementary table 3:** The exclusive 86 bacterial species (OTUs) of the normoglycemic group.

**Supplementary table 4:** The exclusive 15 bacterial species (OTUs) of the impaired glucose tolerance.

**Supplementary table 5:** The exclusive 26 bacterial species (OTUs) of the diabetic group.

**Supplementary table 6:** Shared bacterial species (OTUs) between Normoglycemic and IGT group.

**Supplementary table 7:** Shared Bacterial species (OTUs) between IGT and Diabetic group.

## * List of abbreviations

NGS: Next generation sequencing techniques
16S rRNA: 16S ribosomal RNA gene
Mb: Mega base pairs
BLASTn: Basic Local Alignment Search Tool nucleotide
AGEs: advanced glycation end products
LPS: lipopolysaccharide
TLRs: Toll-like protein receptors
ESRD: end-stage renal disease
OGT: oral glucose tolerance
ADA: The American Diabetes Association
FPG: fasting blood glucose
NHANES: Health and Nutrition Examination Survey
PPD: probing pocket depth
CAL: clinical attachment loss
BOP: bleeding on probing (BOP)
OTUs: operational taxonomic units
HOMD: Human oral microbiome database

## Declarations

### * Ethics approval and consent to participate

This study was approved by institutional review board in King Saud University, Collage of Medicine Riyadh, Kingdom of Saudi Arabia. The subject was provided written informed consent for participating in this study.

### * Consent to publish

All other have consented for publication of this manuscript.

### * Availability of data and materials

All sequence data were submitted to NCBI and submission number is SUB4304342 and BioSample accessions numbers from SAMN09929596 to SAMN09929639 under the BioProject PRJNA488297.

### * Authors’ contributions

**ATMS:** Involved in study conception and design, data analysis and interpretation. Involved in drafting the manuscript or revising it critically for important intellectual content. Preparing the final approval of the version to be published.

**KA:** Involved in study conception and design. Preparing the final approval of the version to be published.

**KD.:** Performed the dental screening and its report preparation.

**BM:** Performed the DNA sequencing and Involved in drafting the manuscript.

**UR:** Involved in acquisition of data, or analysis and interpretation of data Involved in drafting the manuscript.

**MAH:** Involved in study design. Involved in acquisition of data, or analysis and interpretation of data; preparation and involved in drafting the manuscript.

**HT:** Involved in study conception and design. Involved in drafting the manuscript or revising it critically for important intellectual content. Preparing the final approval of the version to be published.

## Acknowledgement

The authors want to thank the members of the University Diabetes Center at King Saud University for their help throughout the study. Special thanks to Dr. Dhekra Alnakeeb and Mrs. Katherine Anne Uy Saeb for their great effort in coordinating the clinical part of the study. In addition, we want to acknowledge that NGS experiments and analysis were supported by the Saudi Human Genome Program (SHGP) at KACST and KFSHRC.

## References

1. IDF Diabetes Atlas [Internet]. [cited 2018 Jan 29]. Available from: https://www.idf.org/e-library/epidemiology-research/diabetes-atlas/13-diabetes-atlas-seventh-edition.html

2. Chee B, Park B, Bartold PM. Periodontitis and type II diabetes: a two-way relationship. Int J Evid Based Healthc. 2013;11:317–29.

3. Bharti P, Katagiri S, Nitta H, Nagasawa T, Kobayashi H, Takeuchi Y, et al. Periodontal treatment with topical antibiotics improves glycemic control in association with elevated serum adiponectin in patients with type 2 diabetes mellitus. Obes Res Clin Pract. 2013;7:e129–38.

4. Moeintaghavi A, Arab HR, Bozorgnia Y, Kianoush K, Alizadeh M. Non-surgical periodontal therapy affects metabolic control in diabetics: a randomized controlled clinical trial. Aust Dent J. 2012;57:31–7.

5. Southerland JH, Taylor GW, Offenbacher S. Diabetes and periodontal infection: Making the connection. Clin Diabetes [Internet]. 2005 [cited 2018 Jan 4];23:171–8. Available from: https://uncch.pure.elsevier.com/en/publications/diabetes-and-periodontal-infection-making-the- connection

6. Kirschning CJ, Wesche H, Merrill Ayres T, Rothe M. Human toll-like receptor 2 confers responsiveness to bacterial lipopolysaccharide. J Exp Med. 1998;188:2091–7.

7. Ahn J, Yang L, Paster BJ, Ganly I, Morris L, Pei Z, et al. Oral Microbiome Profiles: 16S rRNA Pyrosequencing and Microarray Assay Comparison. PLoS ONE [Internet]. 2011 [cited 2018 Jan 4];6. Available from: https://www.ncbi.nlm.nih.gov/pmc/articles/PMC3146496/

8. Docktor MJ, Paster BJ, Abramowicz S, Ingram J, Wang YE, Correll M, et al. Alterations in diversity of the oral microbiome in pediatric inflammatory bowel disease. Inflamm Bowel Dis. 2012;18:935–42.

9. Farrell JJ, Zhang L, Zhou H, Chia D, Elashoff D, Akin D, et al. Variations of oral microbiota are associated with pancreatic diseases including pancreatic cancer. Gut. 2012;61:582–8.

10. Xiao E, Mattos M, Vieira GHA, Chen S, Corrêa JD, Wu Y, et al. Diabetes Enhances IL-17 Expression and Alters the Oral Microbiome to Increase Its Pathogenicity. Cell Host Microbe. 2017;22:120–128.e4.

11. Wade WG. The oral microbiome in health and disease. Pharmacol Res. 2013;69:137–43.

12. Han YW, Wang X. Mobile microbiome: oral bacteria in extra-oral infections and inflammation. J Dent Res. 2013;92:485–91.

13. Peters BA, Wu J, Pei Z, Yang L, Purdue MP, Freedman ND, et al. Oral Microbiome Composition Reflects Prospective Risk for Esophageal Cancers. Cancer Res. 2017;77:6777–87.

14. Zarco MF, Vess TJ, Ginsburg GS. The oral microbiome in health and disease and the potential impact on personalized dental medicine. Oral Dis. 2012;18:109–20.

15. Pizzo G, Guiglia R, Lo Russo L, Campisi G. Dentistry and internal medicine: from the focal infection theory to the periodontal medicine concept. Eur J Intern Med. 2010;21:496–502.

16. Makiura N, Ojima M, Kou Y, Furuta N, Okahashi N, Shizukuishi S, et al. Relationship of Porphyromonas gingivalis with glycemic level in patients with type 2 diabetes following periodontal treatment. Oral Microbiol Immunol. 2008;23:348–51.

17. Aemaimanan P, Amimanan P, Taweechaisupapong S. Quantification of key periodontal pathogens in insulin-dependent type 2 diabetic and non-diabetic patients with generalized chronic periodontitis. Anaerobe. 2013;22:64–8.

18. Thorstensson H, Dahlén G, Hugoson A. Some suspected periodontopathogens and serum antibody response in adult long-duration insulin-dependent diabetics. J Clin Periodontol. 1995;22:449–58.

19. Shillitoe E, Weinstock R, Kim T, Simon H, Planer J, Noonan S, et al. The oral microflora in obesity and type-2 diabetes. J Oral Microbiol. 2012;4.

20. Kampoo K, Teanpaisan R, Ledder RG, McBain AJ. Oral bacterial communities in individuals with type 2 diabetes who live in southern Thailand. Appl Environ Microbiol. 2014;80:662–71.

21. Long J, Cai Q, Steinwandel M, Hargreaves MK, Bordenstein SR, Blot WJ, et al. Association of oral microbiome with type 2 diabetes risk. J Periodontal Res. 2017;52:636–43.

22. Eke PI, Dye BA, Wei L, Slade GD, Thornton-Evans GO, Borgnakke WS, et al. Update on Prevalence of Periodontitis in Adults in the United States: NHANES 2009 to 2012. J Periodontol. 2015;86:611–22.

23. Lang NP, Joss A, Orsanic T, Gusberti FA, Siegrist BE. Bleeding on probing. A predictor for the progression of periodontal disease? J Clin Periodontol. 1986;13:590–6.

24. Hammer O, Harper DAT, Ryan PD. PAST: Paleontological Statistics Software Package for Education and Data Analysis. :9.

25. Kumar S, Stecher G, Tamura K. MEGA7: Molecular Evolutionary Genetics Analysis Version 7.0 for Bigger Datasets. Mol Biol Evol. 2016;33:1870–4.

26. Gomez A, Espinoza JL, Harkins DM, Leong P, Saffery R, Bockmann M, et al. Host Genetic Control of the Oral Microbiome in Health and Disease. Cell Host Microbe. 2017;22:269–278.e3.

27. Yacoubi A. Microbiology of Periodontitis in Diabetic Patients in Oran, Algeria. Ibnosina J Med Biomed Sci [Internet]. 2013 [cited 2018 Jan 4];5:280–7. Available from: http://journals.sfu.ca/ijmbs/index.php/ijmbs/article/view/345

28. Diaz PI, Hoare A, Hong B-Y. Subgingival Microbiome Shifts and Community Dynamics in Periodontal Diseases. J Calif Dent Assoc. 2016;44:421–35.

29. Valentini L, Pinto A, Bourdel-Marchasson I, Ostan R, Brigidi P, Turroni S, et al. Impact of personalized diet and probiotic supplementation on inflammation, nutritional parameters and intestinal microbiota - The “RISTOMED project”: Randomized controlled trial in healthy older people. Clin Nutr Edinb Scotl. 2015;34:593–602.

30. Wang K, Wan Q, Ge L, Yu X-H, Li M, Guo Y, et al. [Preliminary Study on Association Between Intestine Microbiota and Blood Glucose,Blood Lipid Metabolism in Middle-aged and Elderly People in Chengdu]. Sichuan Da Xue Xue Bao Yi Xue Ban. 2018;49:408–13.

31. Conrad R, Ji Y, Noll M, Klose M, Claus P, Enrich-Prast A. Response of the methanogenic microbial communities in Amazonian oxbow lake sediments to desiccation stress. Environ Microbiol. 2014;16:1682–94.

32. Goodson JM, Hartman M-L, Shi P, Hasturk H, Yaskell T, Vargas J, et al. The salivary microbiome is altered in the presence of a high salivary glucose concentration. PloS One. 2017;12:e0170437.

33. Collins MD, Hoyles L, Kalfas S, Sundquist G, Monsen T, Nikolaitchouk N, et al. Characterization of Actinomyces isolates from infected root canals of teeth: description of Actinomyces radicidentis sp. nov. J Clin Microbiol. 2000;38:3399–403.

34. Nakazawa F, Poco SE, Sato M, Ikeda T, Kalfas S, Sundqvist G, et al. Taxonomic characterization of Mogibacterium diversum sp. nov. and Mogibacterium neglectum sp. nov., isolated from human oral cavities. Int J Syst Evol Microbiol. 2002;52:115–22.

35. Chuang L-C, Huang C-S, Ou-Yang L-W, Lin S-Y. Probiotic Lactobacillus paracasei effect on cariogenic bacterial flora. Clin Oral Investig. 2011;15:471–6.

36. Teanpaisan R, Piwat S, Dahlén G. Inhibitory effect of oral Lactobacillus against oral pathogens. Lett Appl Microbiol. 2011;53:452–9.

37. Terai T, Okumura T, Imai S, Nakao M, Yamaji K, Ito M, et al. Screening of Probiotic Candidates in Human Oral Bacteria for the Prevention of Dental Disease. PloS One. 2015;10:e0128657.

38. Abidali H, Sheikh M, Abidali M, Abidali A, Farraji HS, Berry AC. Enterococcus gallinarum Spontaneous Bacterial Peritonitis in an HCV Cirrhotic. Case Rep Hepatol. 2015;2015:898235.

39. Narciso-Schiavon JL, Borgonovo A, Marques PC, Tonon D, Bansho ETO, Maggi DC, et al. Enterococcus casseliflavus and Enterococcus gallinarum as causative agents of spontaneous bacterial peritonitis. Ann Hepatol. 2015;14:270–2.

40. Siqueira JF, Rôças IN. As-yet-uncultivated oral bacteria: breadth and association with oral and extra-oral diseases. J Oral Microbiol. 2013;5.

41. Chen T, Yu W-H, Izard J, Baranova OV, Lakshmanan A, Dewhirst FE. The Human Oral Microbiome Database: a web accessible resource for investigating oral microbe taxonomic and genomic information. Database J Biol Databases Curation. 2010;2010:baq013.

42. Nóbrega LMM, Delboni MG, Martinho FC, Zaia AA, Ferraz CCR, Gomes BPFA. Treponema diversity in root canals with endodontic failure. Eur J Dent. 2013;7:61–8.

43. Fusco V, Quero GM, Cho G-S, Kabisch J, Meske D, Neve H, et al. The genus Weissella: taxonomy, ecology and biotechnological potential. Front Microbiol. 2015;6:155.

44. Klemenak M, Dolinšek J, Langerholc T, Di Gioia D, Micetic-Turk D. Administration of Bifidobacterium breve Decreases the Production of TNF-α in Children with Celiac Disease. Dig Dis Sci. 2015;60:3386–92.

45. Yang C-J, Chang S-Y, Wu B-R, Yang S-P, Liu W-C, Wu P-Y, et al. Unexpectedly high prevalence of Treponema pallidum infection in the oral cavity of human immunodeficiency virus-infected patients with early syphilis who had engaged in unprotected sex practices. Clin Microbiol Infect Off Publ Eur Soc Clin Microbiol Infect Dis. 2015;21:787.e1–7.

46. Hertel M, Matter D, Schmidt-Westhausen AM, Bornstein MM. Oral syphilis: a series of 5 cases. J Oral Maxillofac Surg Off J Am Assoc Oral Maxillofac Surg. 2014;72:338–45.

47. Dewhirst FE, Tamer MA, Ericson RE, Lau CN, Levanos VA, Boches SK, et al. The diversity of periodontal spirochetes by 16S rRNA analysis. Oral Microbiol Immunol. 2000;15:196–202.

48. Byun R, Nadkarni MA, Chhour K-L, Martin FE, Jacques NA, Hunter N. Quantitative analysis of diverse Lactobacillus species present in advanced dental caries. J Clin Microbiol. 2004;42:3128–36.

49. Caufield PW, Schön CN, Saraithong P, Li Y, Argimón S. Oral Lactobacilli and Dental Caries: A Model for Niche Adaptation in Humans. J Dent Res. 2015;94:110S–8S.

50. Verma R, Morrad S, Wirtz JJ. Peptoniphilus asaccharolyticus-associated septic arthritis and osteomyelitis in a woman with osteoarthritis and diabetes mellitus. BMJ Case Rep. 2017;2017.

51. Goldstein EJC, Citron DM, Merriam CV, Warren YA, Tyrrell KL, Fernandez HT. In vitro activity of ceftobiprole against aerobic and anaerobic strains isolated from diabetic foot infections. Antimicrob Agents Chemother. 2006;50:3959–62.

52. Sundqvist G, Reuterving CO. Isolation of Actinomyces israelii from periapical lesion. J Endod. 1980;6:602–6.

53. Honda H, Bankowski MJ, Kajioka EHN, Chokrungvaranon N, Kim W, Gallacher ST. Thoracic vertebral actinomycosis: Actinomyces israelii and Fusobacterium nucleatum. J Clin Microbiol. 2008;46:2009–14.

54. Lee KJ, Kim EJ, Kang SJ, Jang MO, Jang H-C, Jung SI, et al. Lemierre Syndrome Caused by Arcanobacterium haemolyticum Alone in a Healthy Man. Chonnam Med J. 2012;48:190–2.

55. Colombo APV, Boches SK, Cotton SL, Goodson JM, Kent R, Haffajee AD, et al. Comparisons of subgingival microbial profiles of refractory periodontitis, severe periodontitis, and periodontal health using the human oral microbe identification microarray. J Periodontol. 2009;80:1421–32.

56. Paster BJ, Falkler Jr WA, Enwonwu CO, Idigbe EO, Savage KO, Levanos VA, et al. Prevalent bacterial species and novel phylotypes in advanced noma lesions. J Clin Microbiol. 2002;40:2187–91.

57. Fujii R, Saito Y, Tokura Y, Nakagawa K-I, Okuda K, Ishihara K. Characterization of bacterial flora in persistent apical periodontitis lesions. Oral Microbiol Immunol. 2009;24:502–5.

58. Murad CF, Sassone LM, Faveri M, Hirata R, Figueiredo L, Feres M. Microbial diversity in persistent root canal infections investigated by checkerboard DNA-DNA hybridization. J Endod. 2014;40:899–906.

59. Kianoush N, Adler CJ, Nguyen K-AT, Browne GV, Simonian M, Hunter N. Bacterial profile of dentine caries and the impact of pH on bacterial population diversity. PloS One. 2014;9:e92940.

60. Yoo SY, Kim K-J, Lim S-H, Kim K-W, Hwang H-K, Min B-M, et al. First isolation of Streptococcus downei from human dental plaques. FEMS Microbiol Lett. 2005;249:323–6.

61. Douglas GW, Buck LL, Rosen C. Liver Abscess Caused by Haemophilus paraphrohaemolyticus. J Clin Microbiol [Internet]. 1979 [cited 2018 Mar 26];9:299–300. Available from: https://www.ncbi.nlm.nih.gov/pmc/articles/PMC273015/

62. Y S. A Review on the Human Oral Microflora. Res Rev J Dent Sci [Internet]. 2016 [cited 2018 Aug 15];4. Available from: http://www.rroij.com/peer-reviewed/a-review-on-the-human-oral-microflora-78710.html

